# A family of *C. elegans* VASA homologs control Argonaute pathway specificity and promote transgenerational silencing

**DOI:** 10.1101/2022.01.18.476504

**Authors:** Siyuan Dai, Xiaoyin Tang, Lili Li, Takao Ishidate, Ahmet R Ozturk, Hao Chen, Yonghong Yan, Mengqiu Dong, Enzhi Shen, Craig C Mello

## Abstract

Germline Argonautes direct transcriptome surveillance within peri-nuclear membraneless organelles called nuage. In *C. elegans,* a family of Vasa-related Germ Line Helicase (GLH) proteins localize in, and promote the formation of nuage called P granules. Previous studies have implicated GLH proteins in inherited silencing but direct roles in amplification of small RNAs, or in target mRNA or Argonatue binding have not been identified. Here we show that GLH proteins compete with each other to control Argonaute pathway specificity, bind directly to Argonaute-target mRNAs and act to promote the amplification of small RNAs required for transgenerational inheritance. We show that the ATPase cycle of GLH-1 regulates its direct binding to the Argonaute WAGO-1 which engages amplified small RNAs. Our findings support a dynamic and direct role for GLH proteins in inherited silencing beyond their role as structural components of nuage.

## INTRODUCTION

In diverse animals, germline Argonautes direct the transgenerational silencing of transposons and many developmentally important genes (Aravin et al., 2007; Klattenhoff and Theurkauf, 2008; Ozata et al., 2019). In *C. elegans* anti-sense small RNAs (22G-RNAs) produced by cellular RNA-dependent RNA Polymerases (RdRPs) engage WAGO and CSR-1 Argonautes to target nearly all germline-expressed mRNAs. Both RdRP templating and Argonaute surveillance are thought to occur within perinuclear liquid-like condensates called nuage or P granules (Brangwynne et al., 2009; Pitt et al., 2000; Strome and Wood, 1982, 1983; Wolf et al., 1983). Whereas WAGOs mediate silencing, CSR-1 both modulates gene expression (Gurwitz et al., 2016) and protects from silencing (Avgousti et al., 2012; Claycomb et al., 2009; de Albuquerque et al., 2015; Gu et al., 2009; Seth et al., 2013; Tu et al., 2015). The piRNA/PRG-1 pathway engages all transcripts, providing a scanning function that relies on micro-RNA-like base pairing (Shen et al., 2018). PRG-1 recruits RdRP to produce WAGO-associated 22G-RNAs adjacent to PRG-1/piRNA binding sites on more than one-thousand endogenous germline mRNAs, and also promotes small RNA amplification upon encountering foreign RNA sequences (not targeted by CSR-1) (Ashe et al., 2012; Lee et al., 2012; Shirayama et al., 2012). The amplification of WAGO 22G-RNAs and transgenerational silencing are also stimulated by dsRNA through the canonical RNAi pathway, which employs a distinct upstream Argonaute, RDE-1 (Gu et al., 2009; Pak and Fire, 2007; Sijen et al., 2007). WAGO 22G-RNAs, in turn, direct trans-generational gene silencing (Ashe et al., 2012; Buckley et al., 2012; Lee et al., 2012; Shirayama et al., 2012). Although PRG-1/piRNA complexes engage CSR-1 targets (Shen et al., 2018), these interactions—though energetically favorable—fail to stimulate 22G-RNA accumulation and WAGO Argonaute loading in most cases.

DEAD-box proteins have been studied extensively and are known to regulate RNA-RNA and RNA-protein interactions (Linder and Fuller-Pace, 2015). Dead-box proteins are also known to function in a number of Argonaute-small RNA pathways (Pek and Kai, 2011; Wenda et al., 2017; Xiol et al., 2014; Zhang et al., 2012). In insects, for example, VASA functions within nuage to promote the amplification of piRNAs required to suppress transposons (Xiol et al., 2014). In *C. elegans*, the DEAD-box protein RDE-12 functions with the Argonautes RDE-1 and WAGO-1 to promote 22G-RNA production during RNAi (Shirayama et al., 2014; Yang et al., 2014).

Several studies have uncovered roles for a family of <u>G</u>erm <u>L</u>ine DEAD-box <u>H</u>elicase, GLH-family proteins in the regulation of germline RNAs (Beshore et al., 2011; Dallaire et al., 2018). GLH-1 has also been implicated in transgenerational inheritance of RNAi (Spracklin et al., 2017a). Worms with loss-of-function mutations in *glh-1* are viable at 20°C, but become sterile after multiple generations at 25°C, known as a mortal germline phenotype (Kuznicki et al., 2000; Spike et al., 2008a; Spracklin et al., 2017b). *glh-4* null mutant worms exhibit modest fertility defects, but *glh-4 glh-1* double mutants are sterile at all temperatures (Kuznicki et al., 2000; Spike et al., 2008a). Mutations in *glh-1* perturb the localization of other P granule components, including PGL-1 (Chen et al., 2020; Kuznicki et al., 2000; Marnik et al., 2019; Spike et al., 2008a). Several P granule components become completely dispersed in *glh-4 glh-1* double mutants (Chen et al., 2020; Spike et al., 2008a). GLH proteins have intrinsically disordered regions (IDRs), which contain phenylalanine-glycine (FG) repeats that may promote P granule association with nuclear pores (Updike et al., 2011). In addition to the IDR and helicase domains, GLH proteins contain several copies of a retroviral-type (CCHC) zinc finger domain that is also found in the RNA-binding protein LIN-28 (Moss et al., 1997).

Here, we show that GLH-1 and GLH-4 interact with the Argonautes PRG-1 and WAGO-1. We show that GLH-1 and GLH-4 preferentially bind WAGO-target mRNAs as measured by Cross-Linking and IP (CLIP), and this specific binding is diminished in mutants that compromise PRG-1-dependent silencing. We show that GLH-1(K391A), a lesion predicted to disrupt ATP binding by the helicase domain, prevents RNA-duplex unwinding in vitro but does not prevent GLH-1 RNA binding as measured by both *in vivo* CLIP and in vitro gel-shift assays. Indeed, GLH-1(K391A) binding is enhanced not diminished on WAGO target RNAs *in vivo*. A GLH-1 DQAD lesion, predicted to prevent ATP hydrolysis, causes GLH-1 to bind RNA more strongly in vitro. However, interestingly, GLH-1(DQAD) loses its preference for WAGO-target binding in vivo and instead binds many abundant mRNAs including CSR-1 targets.

Our findings suggest that, along with its paralogs, GLH-1 promotes and regulates Argonaute-mediated mRNA surveillance in at least three ways. First, special alleles and double mutants that disrupt P granules also disrupt small RNA levels in the three major nuage-associated Argonaute pathways—i.e., WAGO, CSR-1, and PRG-1—suggesting that GLH proteins help establish a scaffold for Argonaute-mediated surveillance. Second, null alleles of *glh-1* cause a loss of 22G-RNAs on WAGO targets and ectopic 22G-RNAs on many other target mRNAs. The ectopic 22G-RNAs depend on GLH paralogs and PRG-1 activity, suggesting that GLH-1, prevents other GLH paralogs from inducing piRNA-initiated over-production of 22G-RNA on many targets. Third, we show that GLH-1(K391A) strongly impairs transgenerational silencing and WAGO small RNA production without disrupting P granules or other Argonaute small RNA levels. GLH-1(K391A) exhibits a marked increase in its association with the WAGO-1 protein both *in vitro* and *in vivo* suggesting that when unable bind ATP GLH-1 prevents WAGO-1 or co-factors from engaging GLH paralogs, but does so without disrupting GLH protein scaffolding functions required for other Argonaute pathways in nuage.

Our work supports a model in which Argonautes recruit GLH proteins to bind and mark target mRNAs via an ATP-independent mode, that leaves the helicase domain free to bind and remodel RNA duplexes involed in silencing-initiation and amplification. Duplex remodeling by the helicase domain could promote Argonaute release, freeing the Argonaute/guide complex to find new targets, while the GLH protein remains bound to the target RNA through multiple cycles of RdRP transcription, duplex remodeling, and downstream Argonaute loading.

## RESULTS

### GLH-1 and GLH-4 promote piRNA-induced silencing

To identify factors that promote the recruitment of the WAGO silencing machinery by the piRNA pathway, we purified proteins that associate with both the Piwi Argonaute PRG-1 and with the prominent P granule-localized Argonaute WAGO-1. We used CRISPR to engineer the FLAG-coding sequence into the endogenous *prg-1* and *wago-1* genes. We then used FLAG-specific antibodies to recover co-precipitated factors followed by either liquid Chromtographytandem mass spectrometry (LC-MS/MS), or alternatively, by denaturing or native polyacrylamide gel electrophoresis (PAGE) to separate bands containing prominent co-factors for mass spectrometry (IP-MS) (Figure S1A). Using these approaches we identified 32 proteins enriched by FLAG::PRG-1 IP (Figure 1A, Supplementary Table 1A). A similar analysis of the FLAG::WAGO-1 IP identified 126 proteins, including 16 proteins also enriched in the PRG-1 IP (Figure 1A, Supplementary Table 1B).

**Figure 1:**
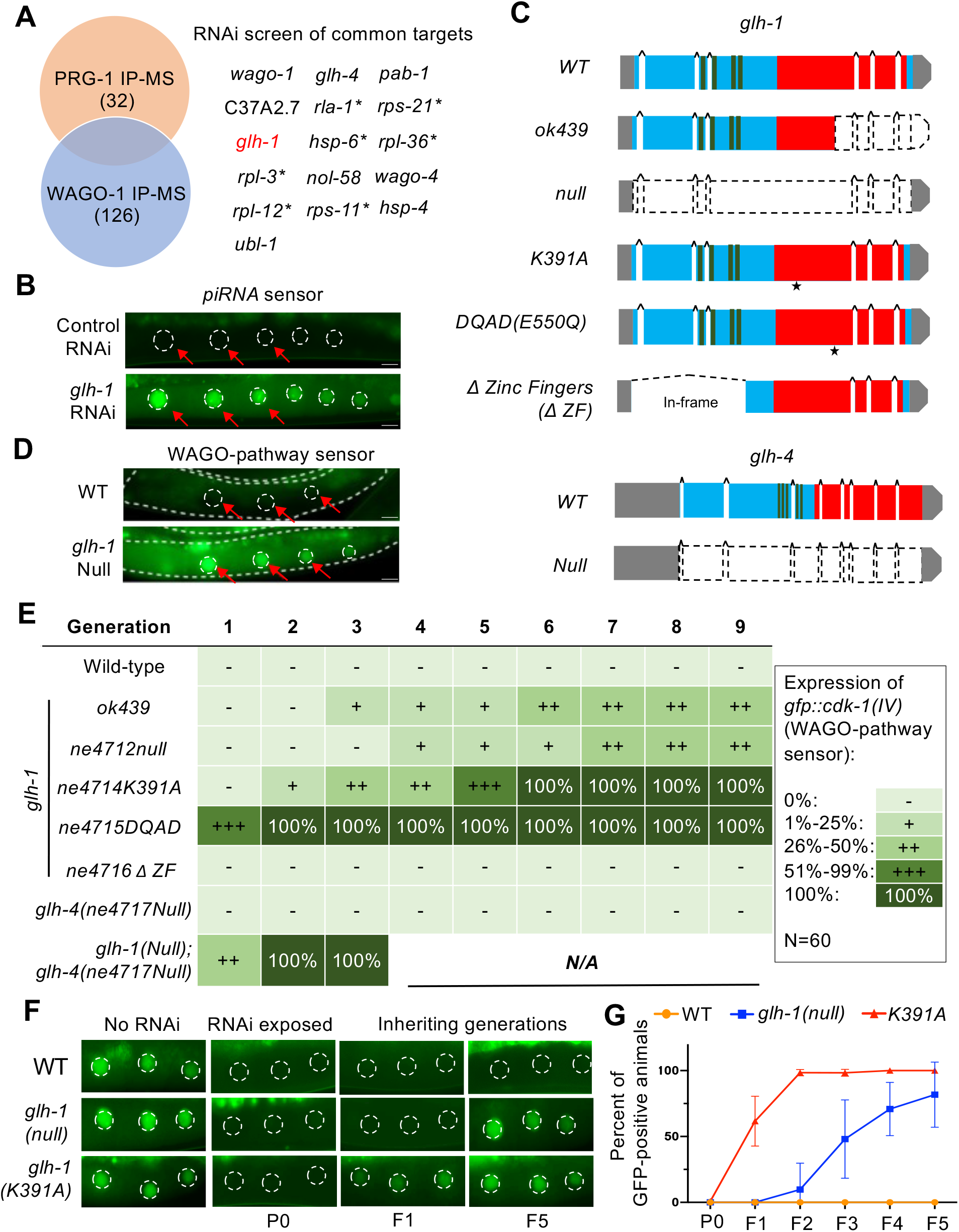
GLH-1 and GLH-4 promote piRNA and RNAi mediated epigenetic silencing. (A): PRG-1 and WAGO-1 IP-MS identified 32 and 126 high-confident cofactors respectively. RNAi screen of 16 candidate genes were performed in a strain with piRNA sensor (*cdk-1::gfp(LGIV);21ux anti-gfp piRNA(X)*). Animals are monitored for at least 10 generations. Genes are essential for *C. elegans* larval development were marked with asterisk. (B): Representative fluorescence images of germline GFP signals showing PIWI-pathway sensor (*cdk-1::gfp(IV);21ux anti-gfp piRNA(X)*) on control RNAi (L4440) and *glh-1* RNAi. Nuclei of oocytes were outlined by dotted lines and were also indicated with red arrowheads. Scale bars represent 10um. (C): Schematic diagram depicting *glh-1* and *glh-4* gene structure and locations of each predicted domain, and mutations in the study. Asterisks indicate amino acid substitution sites. (D): Representative fluorescence images of germline GFP signals showing WAGO-pathway sensor (*gfp::cdk-1(IV))* on control RNAi (L4440) and *glh-1* RNAi. (E) Percentage of worms for WT and each mutant with expressed WAGO-pathway sensors over the generations. Percentage of animals are separated to five categories. (0%: -, 1%-25%: + 26%-50%: ++, 51%-99%: +++, 100%: 100%). At least 60 worms for each generation were scored. (F) Representative images showing expression of *cdk-1::gfp* reporters for worms (WT, glh-1 null and glh-1 K391A) growing on control plates (no RNAi), on *gfp* RNAi plates where dsRNA were present (P0), and followng inheriting generations (1^st^ and 5^th^ generations were shown) upon dsRNA removal. (G) Percent of expressing GFP reporters in WT, glh-1 null and glh-1 K391A worms for each generation. Worms were exposed to *gfp* RNAi (P0) and were taken away from *gfp* dsRNA for following generations (F1 to F5).

Because we wished to find factors that bridge initial targeting by PRG-1 to inheritance maintained by WAGO Argonautes, we asked if knock down of candidates pulled down by both WAGO-1 and PRG-1 affect silencing of two different piRNA pathway reporters: a *cdk-1::gfp* reporter whose silencing depends on an artificial anti-*gfp* piRNA (Shen et al., 2018) (Figure 1B), and a *gfp::cdk-1*, whose silencing was initiated by PRG-1, but is maintained by the WAGO pathway independently of PRG-1 activity (Shirayama et al., 2012).. This analysis identified GLH-1 as a factor required for silencing of both reporters.

Previous studies found that GLH-1 and GLH-4 function together (Kuznicki et al., 2000; Spike et al., 2008a). To investigate if and how GLH-1 and GLH-4 promote piRNA silencing, we generated a series of *glh* mutant alleles that result in complete removal of *glh-1* or *glh-4* coding regions (null), in-frame deletion of the zinc finger domains from GLH-1 (ΔZF), or specific amino acid substitutions in conserved residues of GLH-1 predicted to prevent ATP binding (K391A) or ATP hydrolysis (E550Q; D**E**AD to D**Q**AD) (Figure 1C). Mutations that disrupt ATP binding are known to prevent or reduce RNA binding by DEAD-box proteins (Cheng et al., 2005; Sengoku et al., 2006; Xiol et al., 2014), whereas mutations that disrupt ATP hydrolysis block the release of bound RNA (Hondele et al., 2019; Pause and Sonenberg, 1992; Xiol et al., 2014).

Complete deletion of *glh-1* did not initially de-silence the reporters, but by the fourth homozygous generation, both the piRNA sensor and the WAGO-pathway reporters were expressed in a fraction of *glh-1* null worms, and the percentage of expressing animals increased in subsequent generations (Figure 1D, 1E and data not shown). Interestingly, the K391A and DQAD lesions caused stronger de-silencing phenotypes than did the *glh-1* null alleles (Figure 1E and data not shown). For example, the WAGO-dependent but PRG-1*-*independent *gfp::cdk-1* reporter was de-silenced in 31% of the K391A mutant worms in the second homozygous generation and in 93% of DQAD homozygotes in the first generation. By contrast, in-frame deletion of the CCHC zinc-finger domains in *glh-1(ΔZF)* worms had no effect on silencing (Figure 1E). The onset and extent of silencing defects in these *glh-1* mutants was mirrored by deficits in fertility and embryonic viability (Figure S1B and S1C).

The fact that the GLH-1 K391A and DQAD point mutants have phenotypes that are more severe than the null mutants suggests that they might interfere with the function of paralogs—such as GLH-4—which could otherwise partially compensate for the loss of GLH-1 (Spike et al., 2008a). Indeed, although both reporters remained silent in *glh-4* null worms through nine generations, they were de-silenced in *glh-4(null) glh-1(null)* double mutant worms in the first generation (Figure 1E), a de-silencing phenotype similar to that observed for *glh-1(DQAD)* mutants. These results show that GLH-1 and GLH-4 function redundantly in small RNA silencing, possibly along with PRG-1, but also with the downstream WAGO pathway that propagates piRNA silencing to offspring.

### GLH proteins function in inherited RNAi and prevent cold-sensitive defects in RNAi initiation

Previous studies have shown that *glh-1(null)* mutants have an intact RNAi response in animals exposed to dsRNA but exhibit reduced transmission of silencing to unexposed progeny, an RNAi-inheritance defect (Spracklin et al., 2017b). In agreement with this finding, we found that *glh-1(null)* mutants exhibited complete silencing of a *cdk-1::gfp* reporter when *gfp* dsRNA was present, but upon removal from dsRNA failed to maintain silencing in inheriting generations. For example, whereas wild-type animals transmitted silencing to all progeny in each of the first five generations after dsRNA removal (0% recovery), *glh-1(null)* animals gradually recovered (Figure 1F and 1G). GLH-1(K391A) animals were also fully sensitive to dsRNA silencing in exposed animals, but were more defective than the null in transmission of silencing, reaching nearly 100% de-silencing by the second generation (Figure 1G).

We were surprised to find that presumptive null (deletion or degron) alleles of *glh-1* and *glh-4*—though robustly sensitive to RNAi at room temperature—were partially resistant to RNAi at lower temperatures, a cold-sensitive (cs) RNAi defect (Figure S1D, S1E and S1F). Notably, in contrast to its usually stronger phenotypes, *glh-1(K391A)* worms did not show a csRNAi defect, but instead remained 100% sensitive to RNAi at all temperatures (Figure S1D). Since *glh* null alleles cause ectopic PRG-1-dependent 22G-RNAs (see below), but *glh-1(K391A)* did not, we wondered if misdirection of the PRG-1 pathway might cause the csRNAi defects. Consistent with this idea, a *prg-1* loss-of-function mutation completely suppressed the cold-sensitive RNAi defects of *glh-1* and *glh-4* null mutants (Figure S1E and S1F). Taken together, these findings suggest that the GLH proteins promote inheritance of both dsRNA- and piRNA-induced silencing and also function to prevent a misdirection of the piRNA pathway that causes a cold-sensitive deficit in the dsRNA response (see Discussion).

### GLH-1 lesions affect the localization of P granule components

To correlate the silencing phenotypes of *glh* mutations with their effects on GLH protein localization and P granule integrity, we inserted GFP coding sequences into the endogenous *glh-1* and *glh-4* alleles and analyzed their expression. As expected, both GFP::GLH-1 and GFP::GLH-4 localized to perinuclear P granules (Figure 2A). However, whereas GFP::GLH-1 protein was abundantly expressed throughout the germline, including within the mitotic and meiotic zones, GFP::GLH-4 was expressed primarily within the meiotic region of the germline. (Figure S2A).

**Figure 2:**
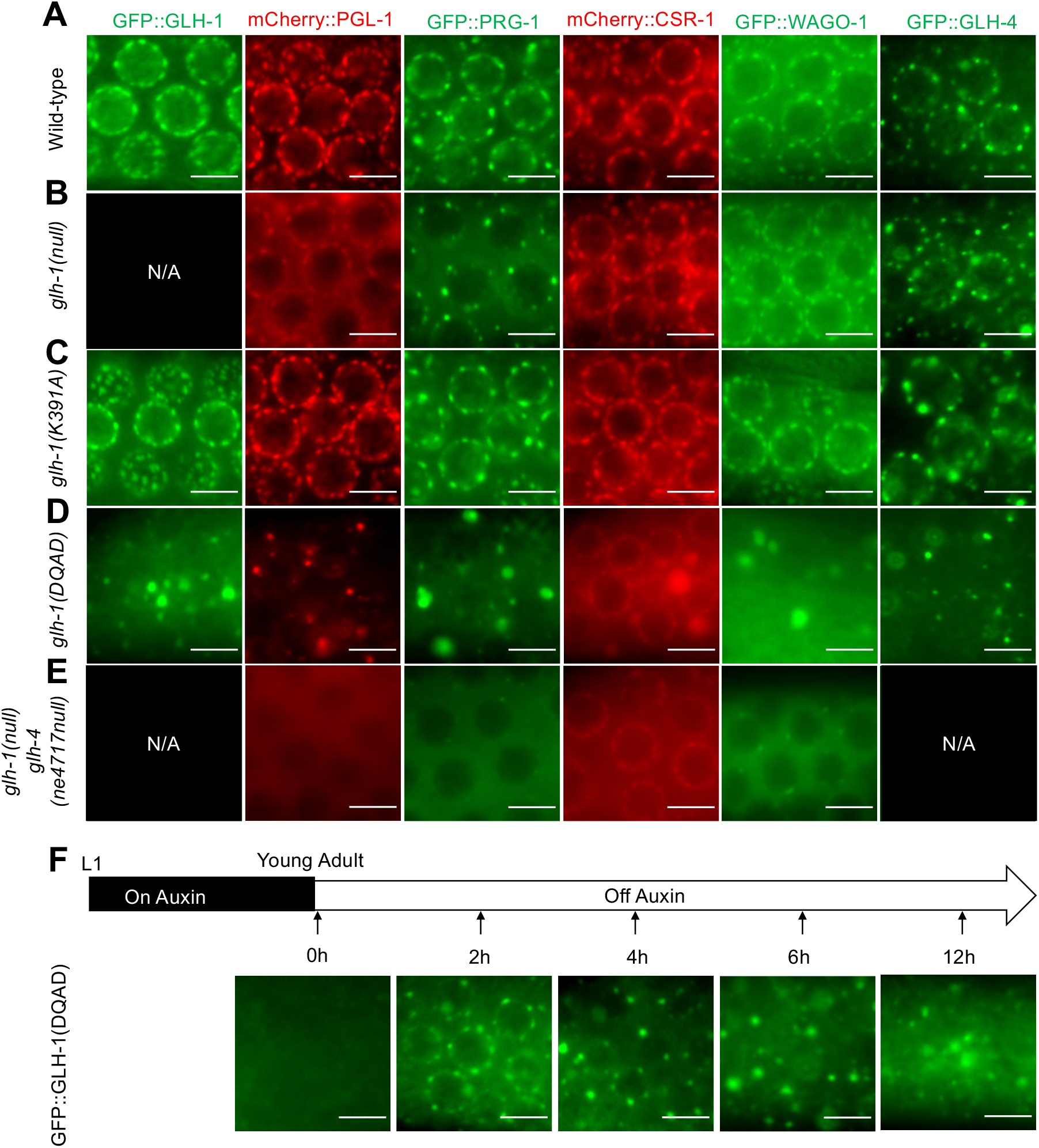
The localization of p-granule components are affected by GLH-1 lesions. A, B, C, D, E): Fluorescence Microscopy showing the sub-cellular localization of P-granule proteins (GFP::GLH-1, mCherry::PGL-1, GFP::PRG-1, mCherry::CSR-1, GFP::WAGO-1 and GFP::GLH-4) in mutant animals with *null* mutations and lesions (K391A, DQAD) introduced to GLH-1 or GLH-4 proteins. Fluorescence proteins (GFP or mCherry) were integrated into endogenous locus of each gene using CSRIPR-cas9 gene editing. Scale bars represent 5um. F)Track the localization of nascent GLH-1(DQAD) protein over the time using Auxin-Induced Degron (AID) system. GFP::GLH-1 proteins with degron tag are depleted by auxin from L1 stage, fluorescent images are captured at indicated time-points after removing animals from auxin. Scale bars represent 5um.

To examine the consequences of individual *glh-1* or *glh-4* mutations on GLH protein localization and on other P granule constituents, we used CRISPR to systematically mutate *glh-1* or *glh-4* in six strains, each expressing a fluorescently tagged P granule component: GFP::GLH-1, GFP::GLH-4, mCherry::PGL-1, GFP::PRG-1, mCherry::CSR-1, or GFP::WAGO-1 (Figure 2). Each CRISPR allele was confirmed by Sanger sequencing and multiple alleles were examined for each lesion. We found that a precise deletion of *glh-1* had mild effects on localization of other P granule components (Figure 2B): PRG-1 and PGL-1 appeared to be slightly reduced in P granules and increased in the cytoplasm, but the localization of GLH-4, CSR-1, and WAGO-1 remained similar to wild type. Interestingly, GLH-1(K391A) was localized to P granules like wild-type GFP::GLH-1, and did not appear to perturb the localization of other P granule components (Figure 2C). As previously reported (Marnik et al., 2019; Chen., et al., 2020) the GLH-1 ATP-hydrolysis mutant, DQAD, dramatically disrupted the localization of P granule components, causing them to disperse in the cytoplasm and to form aggregates in the gonadal rachis (Figure 2D). Although also prominently localized in aggregates, CSR-1 was the only Argonaute that still formed obvious perinuclear foci in the DQAD mutant. In the double *glh-4(null) glh-1(null)*, P granule components formed fewer or, in the case of CSR-1, less pronounced perinuclear foci, and were mainly diffuse throughout the cytoplasm, but did not form aggregates (Figure 2E).

To understand how GLH-1(DQAD) forms aggregates, we controlled the expression of GFP::GLH-1(DQAD) by introducing an in-frame, auxin-inducible degron (Zhang et al., 2015). We maintained the worms on auxin, so that P granules formed properly, and then we removed worms from auxin and followed expression of GFP::GLH-1(DQAD) over time. Initially, the mutant protein localized to perinuclear foci, but within 4 hours of auxin removal, the number of perinuclear GFP::GLH-1(DQAD) foci declined while cytoplasmic aggregates accumulated (Figure 2F). Thus, although GLH-1(DQAD) initially localizes in perinuclear granules that appear wild-type, the continued expression of GLH-1(DQAD) causes the gradual cytoplasmic aggregation of several P granule components over time.

### *glh-1* mutants alter secondary small-RNA levels on mRNA targets

To understand how the GLH proteins regulate small RNA pathways, we used high throughput sequencing to analyze small RNA populations in *glh* mutant worms. Although GLH-1 has been reported to interact with DCR-1 (Beshore et al., 2011), which is required for miRNA biogenesis (Grishok et al., 2001; Hutvagner et al., 2001; Ketting et al., 2001), we did not observe defects in miRNA levels (Fig S3A). Consistent with their stronger defects in silencing, fertility, and embryo viability, GLH-1(K391A) and GLH-1(DQAD) mutants showed the strongest effects on small RNA levels. Many WAGO 22G-RNAs were strongly depleted in the GLH-1(K391A) mutant, whereas piRNAs and CSR-1 22G-RNAs appeared unaffected (Figure 3A and S3C). Expression of a degron GLH-1(DQAD) protein beginning from the L1 stage caused the most severe small RNA defects: piRNAs as well as WAGO and CSR-1 22G-RNAs were all strongly depleted (Figures 3B and S3C). Expressing GLH-1(DQAD) in young adults caused a mild defect during the first several hours after auxin removal, but by 12 hours, piRNA levels were reduced by 70%, WAGO 22G-RNAs by 30%, and CSR-1 22G-RNAs by 10% (Figure S3D). Taken together, these results suggest that GLH-1(K391A) disrupts silencing on WAGO targets, whereas the GLH-1(DQAD) mutant disrupts all the Argonaute small-RNA pathways that localize to P granules, perhaps by causing P granule aggregation (See Discussion).

**Figure 3:**
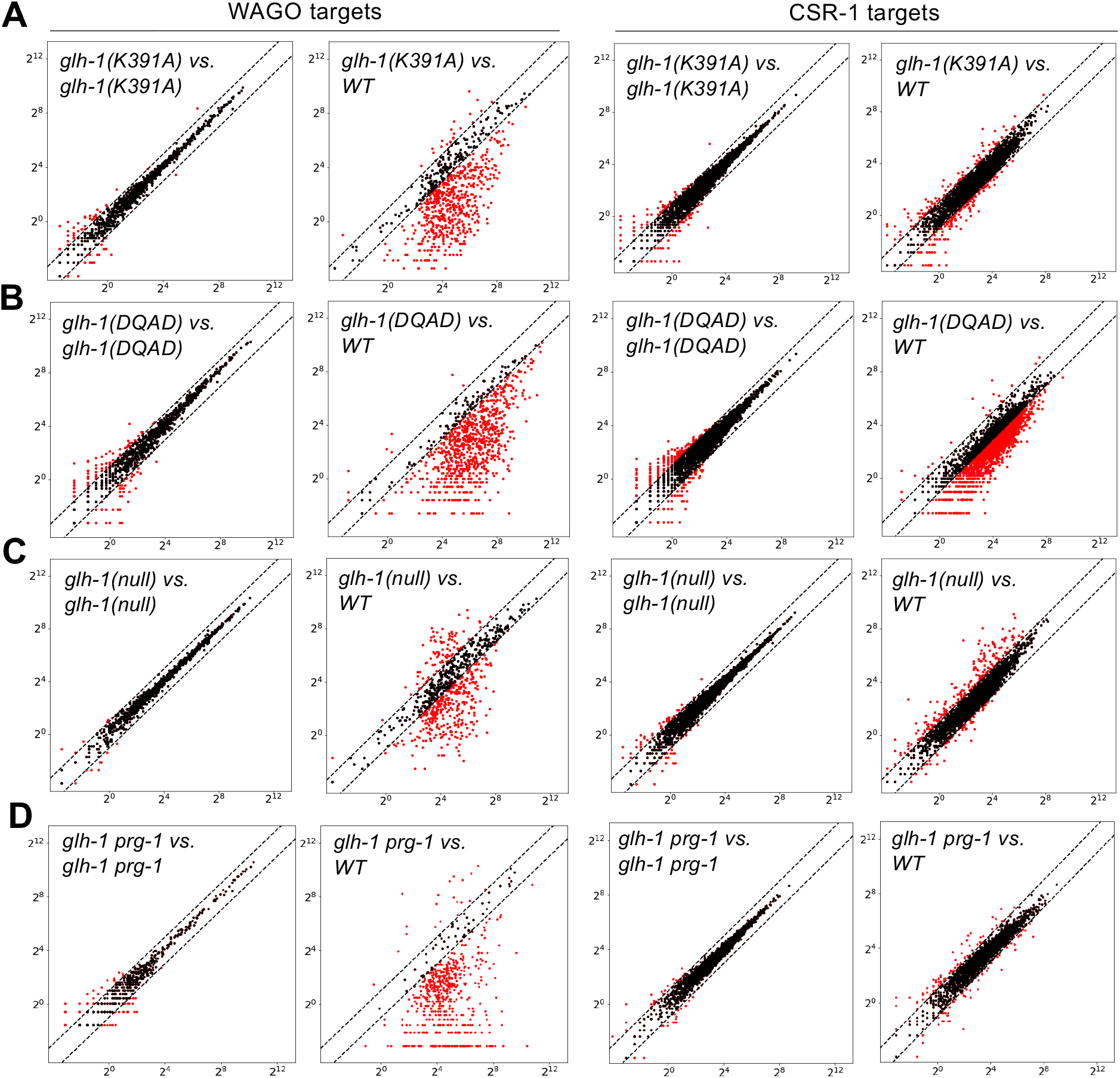
*glh-1* mutants exhibit altered pattern of 22G-RNAs. (A, B, C, D): Scatterplots comparing small-RNA levels for WAGO or CSR-1 targets cloned from wild-type, glh-1(ne4713K391A)(A), glh-1(ne4727DQAD) (B), glh-1(ne4712null) (C) and glh-1(null) in double with degron depleted prg-1 animals. Each dot represents normalized total 22G-RNA reads(reads per million) for a WAGO or CSR-1 target gene. Red dots represent genes that relative small-RNA abundance increase or decrease for at least 2-fold versus wild-type. Dotted lines indicate two-fold threshold.

*glh-1*(*null)* mutants did not alter piRNA levels (Figure S3C) but caused complex changes in CSR-1 and WAGO small RNA levels (Figure 3C). The levels of 22G-RNAs targeting previously annotated CSR-1 targets were increased relative to WT in many cases, while 22G-RNAs targeting previously annotated WAGO targets were both increased and decreased (Figure 3C); these changes were reproducible in the biological replicates (Figure 3C and see below). Although *glh-1 prg-1* double nulls are sterile we were able to conditionally remove PRG-1 from *glh-1(null)* animals using an auxin-inducible *degron::prg-1* allele constructed at the endogenous *prg-1* locus. Exposure to auxin for 24 hours from the L4 stage greately reduced this class of genes with increased 22G-RNA levels for both the WAGO and CSR-1 targets. (Figure 3D).

In order to determine what Argonautes associate with the ectopic 22G-RNAs produced *in glh-1(null)* animals we sequenced small RNAs that co-immunoprecipitated with WAGO-1, WAGO-9, or CSR-1 (Figure 4A). These IP studies confirmed that some WAGO targets made higher levels of WAGO-1 and WAGO-9 bound 22G-RNAs and others made fewer. Strikingly, however, these data revealed that many CSR-1 targets exhibit markedly increased levels of 22G-RNAs that are bound to WAGO Argonautes (Figure 4A). For example, wild-type and *glh-1* null worms produce the same level of CSR-1-bound *mog-4* 22G-RNAs, but *glh-1(null*) worms exhibited a dramatic increase in *mog-4* 22G-RNAs bound to WAGO-1 and WAGO-9 (Figure 4B).

**Figure 4:**
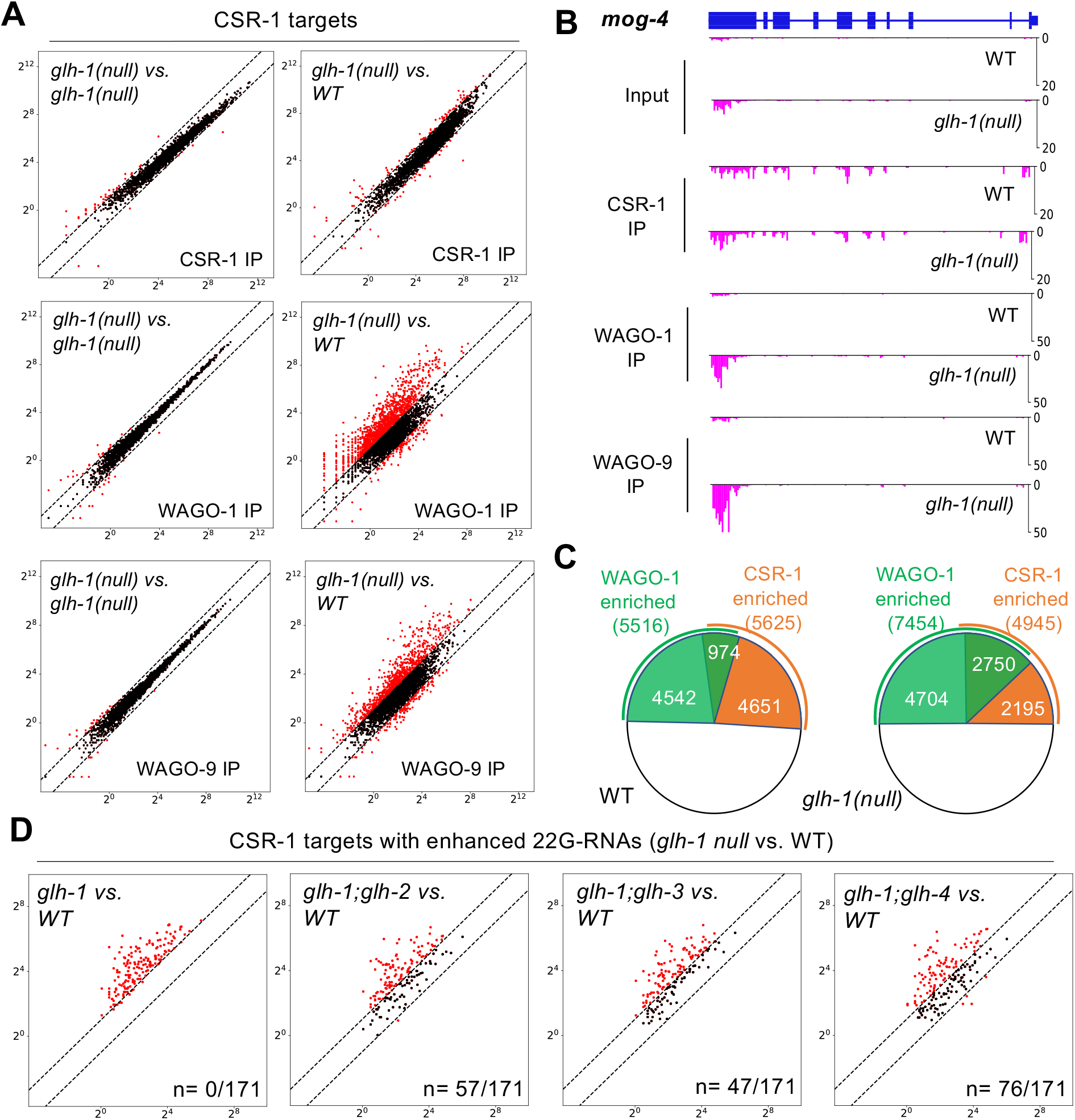
Ectopic WAGO 22G-RNAs in *glh-1 null* animals are enriched over piRNA-targeting sites and depend on GLH paralogs. (A): Scatterplots showing the numbers of small-RNAs recovered from CSR-1, WAGO-1 and WAGO-9 IP for wild-type and *glh-1(null)* worms. (B): Genome browser view of 22G-RNAs aligned to *mog-4* gene that are cloned from CSR-1, WAGO-1 and WAGO-9 IP in wild-type and *glh-1(Null)* animals. Y-axis represents of normalized numbers of 22G-RNAs (reads per million). (C): Piecharts depicting the numbers of genes enriched in CSR-1 and WAGO-1 IP small-RNA sequencing for WT and *glh-1(null)* worms. (D): Distribution of 22G-RNAs that are aligned to piRNA targeting sites in WT and *glh-1(null)* animals. Y-axis represents relative abundance of 22G-RNAs (RPM of 22G-RNA reads/number of C/CLASH density). (E): Scatterplots showing suppression of ectopic 22G-RNAs in degron-depleted *glh-2, glh-3 or glh-4* in double with *glh-1* animals 171 CSR-1 targets with enhanced 22G-RNA levels.

To gain a more comprehensive picture of how *glh-1(nul)l* animals affect Argonaute targeting, we defined sets of genes enriched for binding to the CSR-1 and WAGO-1 Argonautes in our wild-type and mutant IP data sets. We calculated the number of Reads Per Million (RPM) mapping to each gene in the input and IP data sets. A gene was scored as enriched if the RPM level in the IP increased by 2-fold over the level in the corresponding input data set. In wild-type animals, approximately equal numbers of germline genes were enriched in the WAGO-1 and CSR-1 IPs (5516 and 5625 respectively, Figure 4C). Approximately 1000 genes were enriched in both IPs from wild-type worms. Strikingly, in *glh-1(null)* animals the number of WAGO-1-enriched genes increased by 35% to 7454, while the number of CSR-1-enriched genes declined slightly. The percentage of genes enriched in both CSR-1 and WAGO-1 IPs increased from 17% in wild type to 56% in the *glh-1* mutant (Figure 4C). A similar change in targeting was observed for WAGO-9/HRDE-1-associated 22G-RNAs. Thus, in *glh-1* null animals WAGO targeting increases on thousands of germline mRNAs.

### GLH paralogs promote ectopic WAGO 22G-RNA biogenesis

In the absence of GLH-1, other GLH paralogs could promote the production of aberrant 22G-RNAs on target mRNAs. To test this idea, we identified 171 CSR-1 targets showing increased 22G-RNA levels (Supplementary table 2). We then confirmed that the ectopic small RNAs produced on these genes were enriched in WAGO-1 IP or WAGO-9 IP but not in CSR-1 IP (Figure S4C). In order to ask if the ectopic small RNAs depend on GLH-2, GLH-3, or GLH-4, we inserted degron tags into each of these genes in a *glh-1* null mutant strain and then monitored small RNA levels by sequencing after several hours of auxin exposure. In two biological replicates for each strain, we found that auxin-mediated depletion or removal of GLH paralogs suppressed ectopic 22G-RNAs from more than half of the mistargeted genes (106 out of 171; Figure 4D and S4E). For example, exposing *degron::glh-4 glh-1* null double mutants to auxin dramatically reduced ectopic 22G-RNAs at 76 of the 171 mistargeted genes (Figure 4D). Unlike *glh-1(null)* mutants *glh-1(K391A)* did not exhibit ectopic 22G-RNAs on CSR-1 targets (Figure S4D). Taken together, these findings suggest that, in the absence of GLH-1 protein, its paralogs promote the mistargeting, or over amplification, of WAGO 22G-RNAs. GLH-1(K391A) prevents WAGO 22G-RNA induction on piRNA-dependent WAGO targets but also prevents mistargeting perhaps by competing with the GLH paralogs for downstream WAGO 22G-RNA amplification and silencing machinery (see Below).

Multiple-mutants deleting two or more *glh* paralogs are sterile, however we were able to use our small-RNA seq data from the pairwise degron doubles, described above, to create a composite list of genes with GLH-paralog dependent 22G-RNAs. Collectively, summing the genes depleted at least two fold in 22G-RNAs in each double-mutants vs WT identified 2000 genes. The majority of these genes (70%) were among the 1,825 genes depleted 2-fold or more of WAGO 22G-RNAs in *glh-1(K391A),* further supporting the idea that GLH-1(K391A) interferes with the role of GLH paralogs in promoting WAGO 22G-RNA biogenesis (Figure S4F).

### GLH-1(K391A) exhibits enhanced binding to WAGO-1

IP-western blot assays confirmed that GLH-1 and GLH-4 co-precipitate with PRG-1 and WAGO-1 (Figure 5A and 5B). Pretreating the lysate with ribonuclease I (RNase I) reduced but did not eliminate the recovery of GLH proteins in the PRG-1 IP, suggesting that interactions with PRG-1 are only partially bridged by RNA. By contrast, RNase pretreatment greatly reduced the interactions between the GLH proteins and WAGO-1 (Figure 5B), suggesting that these interactions are more strongly dependent on RNA bridging.

**Figure 5:**
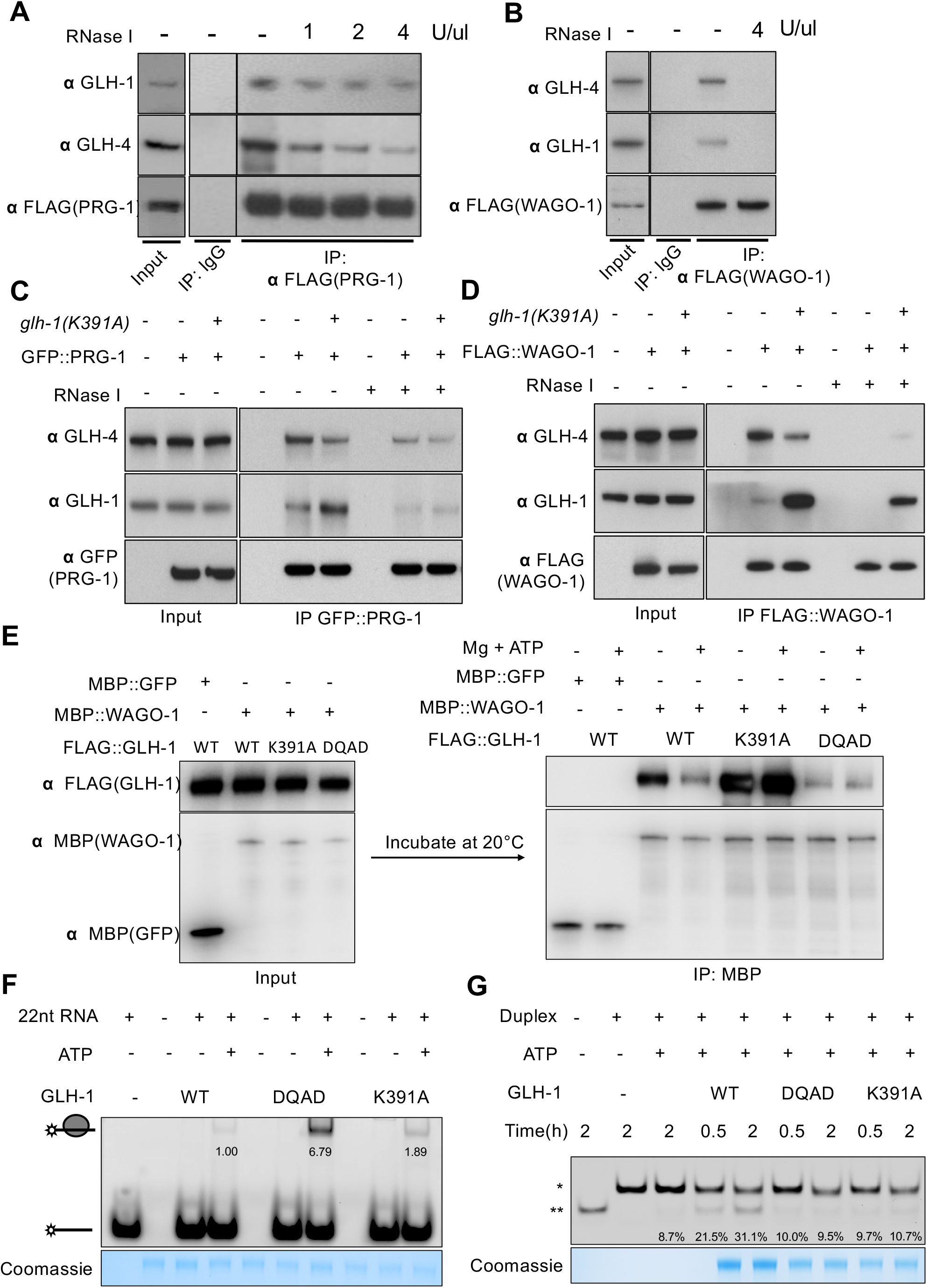
GLH proteins bind PRG-1 and WAGO-1 in a RNA-dependent and independent manner and also affected by ATP cycle. GLH-1 lesions diminish their potent unwinding activities amid binding capabilities are not reduced. (A, B): Co-IP experiment showing physical interactions between WAGO-1 with GLH-1 and GLH-4 (A), between PRG-1 and GLH-1 and GLH-4 (B). Lysates of *flag::prg-1* or *flag::wago-1* worms are incubated with mouse IgG or FLAG-M2 antibodies. Absence (-) or concentration of RNase I treatment are indicated. The blots are probed with GLH-1, GLH-4 and FLAG-M2 antibodies, respectively. (C): Co-IP experiment showing PRG-1 interactions with GLH-1 or GLH-4 in wild-type or mutant animals expressing glh-1(K391A) protein. RNase I treatment were indicated with – or +. PRG-1 were immuno-precipitated by GFP nanobody and eluted fraction were resolved by SDF-PAGE. Blot was probed with anti GLH-1, GLH-4 and GFP antibody. (D): Co-IP experiment showing WAGO-1 interactions with GLH-1 or GLH-4 in wild-type or glh-1(K391A) animals. RNase I treatment were indicated with – or +. PRG-1 were immuno-precipitated by FLAG antibody and eluted fraction were resolved by SDF-PAGE. Blot was probed with anti GLH-1, GLH-4 and FLAG antibody. (E): *In vitro* interactions between MBP::WAGO-1 and FLAG::GLH-1, MBP::WAGO-1 (or MBP::GFP) and FLAG::GLH-1 (or FLAG::GL-1 mutants) were co-expressed in E coli. MBP::WAGO-1/MBP::GFP and their co-factors were purified by MBP affinitry resin, then incubated with Mg2+ and ATP at 20°C for 20 minutes. The purified were resolved by SDS-PAGE following western blots. The blots were probed with anti-FLAG and anti-MBP antibodies. Presence (+) or absence (-) of Mg and ATP were indicated. (F): The binding of synthetic RNA oligos by GLH-1 WT, DQAD and K391A proteins. GLH-1 monomers were expressed in E. coli and purified by anti-FLAG affinity column and Size Exclusion Column. GLH-1 were incubated with 5’ FAM labelled 22nt ssRNA oligos for 30 minutes at RT. The mixture were resolved by native-PAGE and visualized with Bio-Rad imager. (G): In-vitro unwinding assay: Monomeric GLH-1 WT, DQAD and K391A proteins were incubated with RNA duplex with 11-nt 3’ extension for indicated period of time. The products were resolved by native-PAGE, and fluorescence signals were monitored by Bio-Rad imager. RNA duplex were indicated as *, unwound ssRNAs were indicated as **.

To ask if the stronger-than-null phenotypes of GLH-1(K391A) reflect changes in its protein interactions, we first used mass spectrometry to identify proteins that co-precipitate with wild-type GLH-1 or with GLH-1(K391A). This analysis revealed that the K391A lesion does not significantly perturb GLH-1 interactions with other GLH proteins (Figure S5A). We were intrigued to note however, that PRG-1 and WAGO-1 were enriched in GLH-1(K391A) immunoprecipitates (Figure S5A). These findings were confirmed in Co-IPs using GFP-tagged PRG-1 and Flag-tagged WAGO-1, and reciprocal IPs using Flag-tagged GLH-1 and GLH-1(K391A) (Figure 5C, 5D, S5B and S5C). Interestingly, whereas PRG-1 and WAGO-1 interacted more strongly with GLH-4 than with GLH-1, these preferences were reversed in K391A (Figure 5C and 5D). The enhanced interaction between GLH-1(K391A) and PRG-1 was sensitive to RNase treatment. In contrast, the interaction between FLAG::WAGO-1 and GLH-1(K391A) was markedly resistant to RNase treatment (Figure 5C and 5D). We also noted that in GLH-1(K391A) lysates the interaction between WAGO-1 and GLH-4 appeared to be partially resistant to RNase treatment (Figure 5D).

The above findings suggest that ATP binding by GLH-1 regulates a direct (RNA-independent) interaction with WAGO-1. To explore this possibility we performed in vitro binding assays between WAGO-1 and GLH-1 proteins. We tagged WAGO-1 with the Maltose Binding Protein (MBP), and used MBP::GFP as a negative control for specificity. Lysates containing MBP::WAGO-1 or MBP::GFP were mixed with lysates containing wild-type or mutant GLH-1 protein and incubated with or without ATP at 20°C for 20 minutes. MBP pull-down assays showed MBP::WAGO-1 co-precipitates more GLH-1(K391A) than wild-type GLH-1 (Figure 5E), consistent with IPs from worm lysates. Adding magnesium and ATP prior to MBP-pull-down slightly reduced WAGO-1 binding to wild-type GLH-1, but did not affect binding to GLH-1(K391A) or GLH-1(DQAD) (Figure 5E). The control MBP::GFP fusion protein did not bind to GLH-1 (Figure 5E). These data suggest that WAGO-1 preferentially interacts with GLH-1 when the helicase domain is in an ATP-unbound conformation.

### GLH-1 has RNA binding and unwinding activities *in vitro*

We next asked whether GLH-1 can bind RNA and unwind RNA duplexes *in vitro*. To check for RNA binding, we incubated wild-type or mutant GLH-1 proteins with fluorescently labeled 22-nt single-stranded (ss)RNA oligo and visualized any resulting protein/RNA complexes as gel shifts by native PAGE. We found that both GLH-1 WT and K391A showed weak binding to the ssRNA oligo, whereas GLH-1(DQAD) showed strong ATP-dependent binding (Figure 5F).

To examine unwinding activity, we incubated the recombinant GLH-1 proteins with Mg^2+^ and ATP and a short fluorescently labeled duplex RNA containing an 11-nt 3’ extension. In the presence of excess single-stranded competitor lacking the 11-nt extension, the unwound labeled molecules will anneal with the competitor to produce a labeled product that can be distinguished from the substrate by PAGE. Indeed, this shorter labeled product accumulated over time in the presence of wild-type GLH-1 protein (Figure 5G). As expected, unwinding required ATP and was reduced by the K391A and DQAD mutations (Figure 5G).

### GLH-1 and GLH-4 associate with overlapping sets of WAGO target mRNAs

To determine if GLH proteins bind RNA *in vivo*, we used a modified cross-linking and immunoprecipitation (CLIP) assay (Chi et al., 2009, Norstrand et al., 2016) to analyze four proteins—GLH-1(WT), GLH-1(K391A), DEGRON::GLH-1(DQAD) and GLH-4(WT), all tagged with the FLAG epitope at the endogenous locus. In order to measure background binding to the matrix, adult worms expressing each FLAG-tagged GLH-1 proteins were analyzed in parallel with control strains expressing the corresponding untagged *glh-1* alleles. Populations of DEGRON::GLH-1(DQAD) worms were grown to adulthood in the presence of auxin to deplete the toxic protein, and then auxin was removed to allow GLH-1(DQAD) to accumulate in P granules for either 2 or 4 hours before crosslinking.

Using a 2-fold enrichment over background binding, CLIP-seq analyses identified 3120 GLH-1(WT), 3927 GLH-1(K391A), 4439 GLH-1(DQAD), and 6158 GLH-4 targets (Figure 6A). The majority of RNA fragments associated with GLH-1(WT), GLH-1(K391A), GLH-1(DQAD) or GLH-4 (88%, 88%, 79% and 58% respectively) derived from mRNAs (Figure S6A). piRNAs were also recovered at high levels, especially in GLH-4 IP complexes, where they made up 25% of the reads; by comparison piRNAs comprised 5%, 3%, and 1% of reads in GLH-1(WT), GLH-1(K391A) and GLH-1(DQAD) complexes (Figure S6A). Although we expected K391A to disrupt RNA binding, we were surprised to find that K391A CLIP enriched nearly 30% more mRNAs than WT (Figures 6A and 6B). Moreover, GLH-1(K391A) enriched 89% of the targets enriched by GLH-1(WT) (Figure 6B).

**Figure 6:**
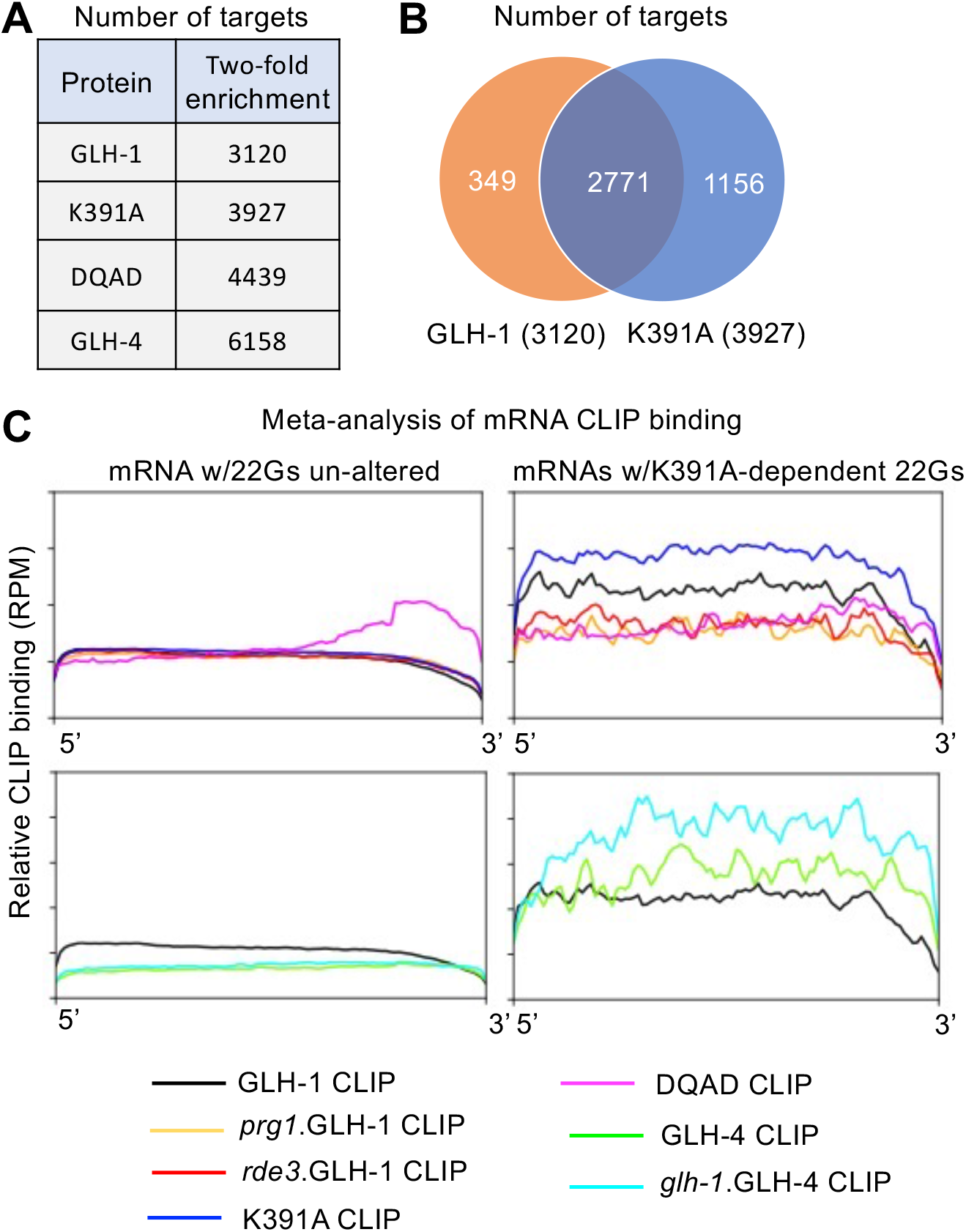
GLH-1 and GLH-4 bind predominantly WAGO target mRNAs whose 22G-RNAs are reduced in glh-1 K391A animals. (A): Table depicting the numbers of genes with at-least two-fold enriched binding of indicated proteins. (GLH-1 WT, GLH-1 K391A, GLH-1 DQAD and GLH-4) (B): Venn Diagram representing counts of genes that are targeted by GLH-1 (Orange circle), GLH-1 K391A (Blue circle) and both proteins at 2-fold enrichment. (C): Metagene analysis of GLH-1 WT (black), GLH-1 K391A (blue) and GLH-1 DQAD (megenta), also GLH-1 WT in *prg-1* (orange) and *rde-3* (red) mutants on control category and GLH-1 K391A depletion category. K391A depletion category was defined by genes showing at-least two-fold reduction in 22G-RNA levels and more than 5 RPM in *glh-1(K391A)* mutants compared to WT animals. Y-axis indicates averaged CLIP binding reads along each mRNA as a percentage of the mRNA length and plotted the aggregate binding to each interval extending from 0% (5’) to 100% (3’) along the length of the transcript. Metagene analysis of GLH-1 WT (black), GLH-4 WT (green) and GLH-4 in *glh-1* mutants (Torquiose) on control category and GLH-1 K391A depletion category.

**Figure 7:**
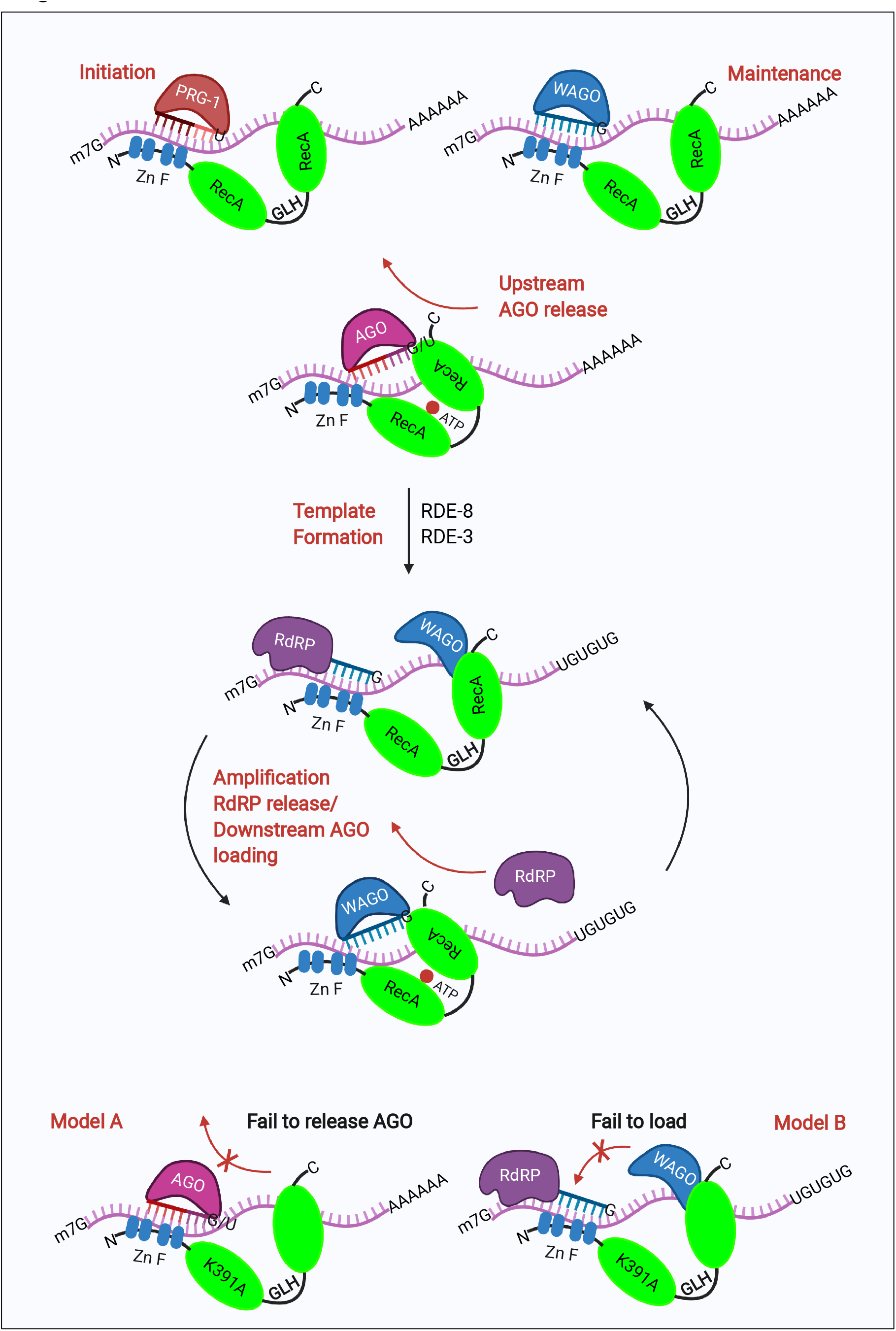
GLHs control the speficity of Argonaute pathways and drive small-RNA amplification cycle by promoting duplex remodeling and WAGO loading. GLHs were recruited to the transcripts in response to PRG-1 pathway for silencing initiation and WAGO pathways for silencing maintenance. Upon target recongnition, the transcripts were processed by RDE-8 and RDE-3 to generate polyUG tailed template RNAs for anti-sense 22G-RNA amplification. GLHs promote the cycles of 22G-RNA production by releasing the Argonautes from the transcripts with their helicase activities for RdRP recruitment. Additionally, GLHs associates with empty WAGO Argonautes in ATP-dependent manner to template RNAs for WAGO loading to nascent 22G-RNAs. Retention of GLHs to target site by helicase-independent activities enable multiple rounds of amplification.

To explore the relationship between GLH mRNA binding and 22G-RNA levels, we analyzed the subset of mRNAs whose 22G-RNA levels decreased by at least 2-fold in GLH-1(K391A) mutants. We chose this group of targets because the K391A mutant strongly depletes a subset of PRG-1-induced WAGO 22G-RNAs that depend on GLH-1 and/or its paralogs (i.e., GLH-dependent targets) without perturbing P granule scaffolding functions required for other germline Argonaute pathways. We used the remainder of germline mRNAs as a control set. To reveal how CLIP binding distributes along the length of mRNAs, we performed a meta-analysis of GLH binding along GLH-dependent or control mRNAs (see STAR Methods). As expected, wild-type GLH-1 was enriched on GLH-dependent targets but not on the control set (Figure 6C: top panel). Moreover, enriched binding required PRG-1 and RDE-3 activities, consistent with the idea that direct binding of wild-type GLH-1 correlates with GLH-dependent 22G-RNA production on these targets. Compared to wild-type GLH-1, GLH-1(K391A) showed enhanced binding along these same targets (Figure 6C: top panel).

GLH-1(DQAD), in contrast, showed a strikingly different pattern of mRNA binding. At 2 hr after auxin removal, when GLH-1(DQAD) is properly localized to intact P granules (Figure 2F), DQAD primarily enriched mRNAs (79% of CLIP reads). Notably, however, the mRNAs enriched by DQAD were not restricted to GLH-dependent targets, but also included many GLH-1-independent targets. Moreover, DQAD preferred to bind the 3’ regions of these mRNAs (Figure 6C: top panel), including many CSR-1 targets.

Consistent with competition between GLH-1 and GLH-4 for WAGO 22G-RNA induction, GLH-4 binding to GLH-1-dependent target mRNAs was increased in a *glh-1(null)* mutant (Figure 6C: bottom panel). In a separate meta-analysis, we found that sites of ectopic 22G-RNAs in *glh-1* mutants correlate with ectopic or increased GLH-4 binding (Figure S6B). Taken together these findings suggest that GLH-1 and GLH-4 binding to target RNAs correlates with the production of PRG-1 and RDE-3-dependent small RNAs.

## DISCUSSION

### A link between the GLH-1 ATP cycle and Argonaute surveillance

A poorly understood aspect of transgenerational silencing is how targeting by a primary Argonaute (PRG-1 for piRNAs and RDE-1 for the dsRNA response) leads to the amplification of secondary small RNAs that are transmitted to offspring. Here, we have shown that members of an expanded family of DEAD box proteins related to Drosophila VASA, (the GLH proteins) physically interact with Argonaute proteins and with target RNA to promote transgenerational silencing. RNA-binding by the helicase domain of DEAD-box proteins is gated by ATP-binding, while release requires ATP hydrolysis (Yang et al., 2005; Liu et al., 2008; Xiol et al., 2014). Surprisingly, a lesion expected to prevent ATP binding, GLH-1(K391A), failed to prevent *in vitro* RNA binding, and instead caused enhanced association with Argonaute-pathway target mRNAs *in vivo*. A glutamate to aspartic acid lesion in the DEAD box (motif II), GLH-1(DQAD), exhibited enhanced *in vitro* binding (as expected), and exhibited Argonuate-pathway-non-specific association with the 3’ ends of mRNAs *in vivo*.

Taken together our findings are consistent with a model the PRG-1 and WAGO-1 pathways recruit GLH-1 to bind specific targets via a Lysine 391-independent (and hence likely a helicase-independent) RNA-binding activity. Positioning of a GLH-1 complex on target mRNA via a helicase-independent binding activity could enable the piRNA and WAGO Argonaute systems to employ GLH proteins to promote cycles of RNA-duplex remodeling required for inherited WAGO silencing. Consistent with this idea, GLH-1(K391A) exhibited enhanced binding on piRNA and WAGO target mRNAs and to PRG-1 and WAGO-1 proteins. Perhaps when locked in the open conformation, the helicase domain of GLH-1(K391A) is unable to promote PRG-1 or WAGO-1 release from target mRNA, causing normally transient initiation complexes to accumulate on target mRNA (see Model Figure A). Alternatively, or additionally, in the absence of ATP binding, inefficient unwinding of nascent 22G-RNA/template RNA duplexes generated by RdRP, might stabilize an interaction between GLH-1(K391A) and newly expressed, unloaded WAGO, thereby trapping a normally transient amplification complex (a GLH-1-WAGO-1 loading complex, see Model Figure B). Like other DEAD-box proteins (Yang et al., 2005), GLH-1 can bind and remodel short RNA duplexes upon ATP hydrolysis. Moreover, in its ATP unbound form GLH-1 preferentially associates with WAGO-1 a downstream Argonaute involved in amplification. Thus it is possible that the ATPase cycle of GLH-1 directly couples release of the upstream Argonaute to the arrival of an unloaded WAGO required for amplification (Model). Retention of the helicase at the target site by a helicase-independent RNA binding modality could allow the upstream Argonaute to leave while enabling the GLH protein remains bound enabling multiple rounds of amplification. In insects and mammals, Piwi Argonautes amplify epigenetic signals on transposon RNAs via an amplification cycle in which the cut site of a loaded Argonaute creates the 5’ end of a guide RNA that is loaded onto a downstream Argonaute. This downstream Argonaute can then cleave a corresponding antisense transcript to regenerate the upstream guide, and so on. Interestingly, this “ping-pong” amplification cycle requires the DEAD-box protein VASA, and studies in insect cells have linked VASA ATP binding with formation of a Piwi complex, called the “amplifier complex” (Xiol et al., 2014). A VASA-K230N mutation analogous to GLH-1(K391A) disrupts binding to the PIWI orthologs (Siwi and AGO-3) that mediate the ping-pong cycle. In contrast, the VASA(DQAD) mutant protein associated with both Piwi Argonautes, albeit bridged by RNA. Interestingly, this VASA(DQAD) complex contained Siwi-associated piRNAs but not AGO3-associated piRNAs, suggesting that the mutant VASA(DQAD) protein traps an intermediary complex in the ping-pong cycle, prior to loading the downstream AGO3 Argonaute. Although the details differ, these findings and those from the present study hint at a conserved role for the ATPase activities of VASA and GLH-1 in the amplification of small RNA silencing.

## Conclusion

Here we have shown that the VASA homologs GLH-1 and GLH-4, which are required for P granule assembly and homeostasis (herein; Chen et al., 2020; Spike et al., 2008a), play complex and dynamic roles in Argonaute surveillance. Although the GLH paralogs have often been considered redundant factors, our studies reveal major differences in their mRNA, small-RNA, and protein interactions. For example, our analysis suggests that GLH-1 competes with or can prevent recruitment of GLH paralogs on many target mRNAs. Thus, mutating one GLH factor causes mistargeting of the other. Mistargeting of GLH paralogs in turn requires PRG-1 and results in expression of ectopic WAGO pathway small RNAs on thousands of germline mRNAs, including many CSR-1 pathway targets. Mistargeting of the WAGO pathway to thousands of additional mRNA s likely explains the cold-sensitive defect of *glh-1* null mutants, as ectopic piRNA-dependent WAGO silencing could reduce availability of WAGO machinery to function in RNAi-mediated silencing. Alternatively, piRNA-dependent mistargeting could silence one or more genes that encode protein effectors of WAGO silencing, as suggested in the recent studies by Ouyang et al., (2019) and Dodson and Kennedy (2019). Our findings reveal nuage as a complex organelle where RNA-binding factors interact to impose delicate regulation on mRNA homeostasis and expression. Understanding the cascading effects of mutations that shift the balance of RNA binding and surveillance in nuage could shed light on related perturbations in RNA-binding factors that cause a myriad of human disorders.

## Materials and Methods

### Immunoprecipitation and Mass spectrometry

100,000 synchronized *flag::tev::prg-1 and flag::tev::wago-1* worms were homogenized by FastPrep-24 (MP Biomedicals) in lysis buffer (20mM HEPES PH=7.0, 250mM Sodium Citrate, 0.5% Triton X-100, 0.1% Tween-20, 1mM DTT). The same experimental procedures were also performed for N2 worms as negative controls. Protein extracts were incubated with 3ul anti-FLAG antibody (Clone M2, Sigma-Aldrich) and 50ul Protein-G magnetic beads (Thermo Scientific) at 4 Celsius for 2 hours. Prior to elution, pull-downed components were pretreated with RNase for removing proteins bridged by RNAs. Immunoprecipitates were released from beads by TEV protease cleavage (Thermo Scientific). Finally, the purified protein complex were resolved by SDS-PAGE and visualized by Silver staining (Pierce). The elutes were precipitated by acetone and air dried. Protein samples were re-suspended and digested with Trypsin. LC-MS/MS were conducted at Mengqiu Dong’s lab in NIBS. MS data were processed as described previously (Feng et al., 2017). Proteins with at-least two fold enrichment of relative peptide levels in IP samples over negative control were identified as co-factor of PRG-1 and WAGO-1. See Supplementary table 1 for proteins and their descriptions.

### RNAi Screen

RNAi screen were conducted as previously described. In brief, worms were fed with HCT115 E. coli expressing either control dsRNA (L4440) or dsRNA targeting 16 desired genes for more than 10 generations. All worms were growing at 20°C and at least 60 animals were scored per generation. RNAi that causes larval lethality was indicated in figure 1.

### Generation of worms by CRISPR

C elegans strains were produced by CRISPR-cas9 RNP methods (<u>Dokshin et al., 2018</u>, Ghanta et al., 2021) by co-injection Cas-9 RNPs and rol-6 marker. At-least two independent alleles were generated for each strain.

### Desilencing of transgene reporters in C. elegans

Published glh-1 alleles and other lesions on other *glh-1 and glh-4* genes were introduced to *piRNA sensor, cdk-1::gfp;21ux anti-gfp LGIV* and wago pathway sensor *gfp::cdk-1 LGIV* worms by crossing and by CRSIPR-cas9 method. All worms were grown at 20°C and at least 60 worms were scored by fluorescence microscopes. Animals were passaged by random picking 5-6 worms to new plates for producing progenies of next generation.

### Assays for assessment of RNAi inheritance and cold-sensitive RNAi

For inheritable RNAi assay, synchronous population of worms were grown on *gfp* RNAi or L4440 RNAi for one generation, then the worms were bleached and plated progenies on OP50 food. Fluorescent signals of 100 worms were checked for each generation. For cold-sensitive RNAi assay, all worms were grown on *pos-1* RNAi or L4440 RNAi food and maintained at 15°C, 20°C and 25°C respectively until they laid sufficient eggs for scoring.

### Fertility of *C. elegans*

Synchronized populations were bleached and grown at 20°C. Brood sizes and hatching rates were measured by counting eggs laid onto the plates and percent of viable embryos. Progenies of at least 15 animals were scored for each strain.

### Microscopic Imaging of P granule components

Images of GFP-tagged or mCherry-tagged P granule components were captured by Zeiss M2 microscopes. Images were processed and cropped by ImageJ. GFP::AID::GLH-1(DQAD) worms were kept on the plates with 500mM IAA (Afar aesar) until they grow to young adults. Worms were then removed from Auxin plates and allow *de novo* expression of GLH-1(DQAD) over the time for imaging.

### Preparation of small-RNA libraries and deep sequencing

Total RNAs were extracted by synchronous population of worms by Trizol(Simga Alrich) and isopropanol precipitation. Small-RNAs were further purified by mir-Vana miRNA isolation kit (Thermo Scientific). Small-RNA cloning was carried out as previously described (Shen et al., 2018). Briefly, homemade PIR-1 were used for remove 5’ triphosphate of endogenous 22G-RNAs. The 3’ and 5’ adaptors were ligated by truncated ligase 2 (NEB) and ligase 1 (NEB) sequentially. cDNA were produced by superscript III reverse transcriptase (Thermo). cDNAs were amplified by Q5 polymerase and barcodes sequences were added. Amplified products were subjected to Native PAGE and size selection. High throughput sequencing was conducted using Illumina High-Seq, Next-Seq and Nova-Seq platforms . For sequencing of Argonaute-bound small-RNAs, Argonaute proteins/small RNA complex were extracted from 50,000 synchronized worms and immunoprecipitated with anti-flag antibody (Sigma Aldrich) or GFP nanobody (Thermo). Small-RNAs were then extraced by Trizol and purified using ethanol precipitation. Library preparation approach remains the same as the above mentioned.

### Analysis of small-RNA sequencing

Raw sequencing data were demultiplexed, 3’ adaptor trimmed and aligned to WS262 and WS272. Unique mapped 22nt-long anti-sense reads were used for later analysis. Relative numbers of reads mapped to each gene were normalized by total read counts of each sample (Reads per million, RPM). Genes enriched in IP WAGO-1 and CSR-1 were scored by limiting minimal 1 RPM and more than 2-fold enrichment over the level in corresponding input data. Figures were generated using Excel, Prism 8.0, Pandas, Matplotlib and Deeptools.

### Co-immunoprecipitation and western blotting

50,000 Animals were homogenized in lysis buffer and their lysates were incubated with 2ul FLAG antibody (Clone M2, Sigma) together with 30ul Protein G beads for FLAG-tagged WAGO-1 IP, or GFP nanobody (Thermo) for GFP-tagged PRG-1 IP. Lysates were per-treated with RNase I. Immunoprecipitates were boiled with 4X SDS sample buffer (BioRad) and were resolved by SDS-PAGE later transfer to PVDF membranes. GLH-1 antibody (1:5000), GLH-4 antibody (1:2000), anti-FLAG Hrp-conjugated (Sigma, 1:5000), anti-V5 antibody were used for probing the blots.

### In-vitro protein binding assay

MBP::WAGO-1, MBP::GFP, GLH-1 WT and its variants were constructed onto pET-Duet1 vectors. FLAG-tagged GLH-1 WT, FLAG-tagged GLH-1 variants, MBP-tagged GFP and MBP-tagged WAGO-1 proteins were expressed in BL21 cells following 0.5mM IPTG treatment for 18 hours at 18 C. *E. coli* were collected and resuspend them in lysis buffer (50mM Tris-HCl pH 8.0, 150mM NaCl, 1mM PMSF, 2mM DTT). Bacterial homogenates were then sonicated and went through MBP-trap affinity column by AKTA. With stringent washing with lysis buffer, protein complex were eluted with 10mM Maltose (Sigma Aldrich). Resolve elutes by SDS-PAGE and transfer to PVDF membrane for western blots. FLAG-hrp (Abcolonal, 1:5000) and MBP antibody(MBL, 1:2500) were used for probing the blots.

### Protein purification

The full-length FLAG-tagged GLH-1 was cloned into pET28a vector. The point mutations GLH-1 K391A and GLH-1 E500Q were introduced by mutagenesis. All constructs were expressed in ER2566 E.coli cells (WEIDI) at OD600 of 0.8 with 0.5mM IPTG for 18h at 18℃. Cells were resuspended in lysis buffer (50mM Tris-HCl pH 8.0, 500mM NaCl, 1mM PMSF, 2mM DTT) and was sonicated on the ice. Then the lysates were treated with 10μL Ambion® RNase Cocktail™ (Invitrogen) and 10μL Micrococcal Nuclease (New England Biolabs) for removing bacterial RNAs that remain bound to the proteins. After high-speed centrifugation, the supernatant was loaded onto a column with 1mL Anti-DYKDDDDK G1 Affinity Resin (GenScript) and the 1×FLag-tagged proteins were affinity extracted. Then the resin was washed with 60mL wash buffer (50mM Tris-HCl pH8.0, 500mM NaCl) and the proteins were eluted with 2mL elution buffer (50mM Tris-HCl pH8.0, 500mM NaCl, 0.8mg/mL flag peptide). After affinity purification, eluted samples were run on Superdex 200 in gel filtration buffer (25mM Tris-HCl pH8.0, 500mM NaCl, 2mM DTT) to separate non-functional aggregates and functional monomers according to their molecular weight difference. All chromatography was conducted at 4℃. The abundance and purity of each collected fraction was detected by Coomassie blue staining on SDS-PAGE. Finally, the fractions with clean monomers were collected, and their concentration were quantified by BCA Protein Assay Kit (Beyotime) for further use.

### Electrophoretic Mobility Shift Assay (EMSA)

RNA binding activities of purified GLH-1 WT and mutants K391A and E500Q were assessed by incubating 10μM monomeric proteins and 2μM 5’ end Cyanine-3 (Cy3) labelled RNAs (22nt RNA 5’-Cy3-GUCAAAGAUAGCCUUGACCUUG-3’ for GLH-1 WT, mutants K391A and E500Q) in the presence and absence of 2mM ATP for 30min at RT in binding buffer (25mM HEPES pH7.4, 150mM NaCl, 2mM MgCl2). The RNA-proteins complexes were separated on native 8% polyacrylamide gel at 60V in 1×TBE buffer in cold room. Fluorescence signals were detected by BioRad imager with a proper filter.

### In-vitro helicase unwinding assays

RNA duplex with 3’ overhangs were prepared by annealing 5’ Cyanine-3 labelled 11nt RNA (5’-Cy3-AGCGCAGUACC-3’) with a 22nt RNA (5’-GGUACUGCGCUUUUAUGACAUC-3’). 20μM purified GLH-1 proteins (WT, K391A or DQAD) was incubated with 25nM RNA duplex in helicase buffer (25mM HEPES pH7.4, 150mM NaCl, 2mM MgCl_2_, 2mM ATP) over the time. Unlabeled 11nt RNA was added to final concentration of 0.125μM to prevent re-annealing of unwound products. The reactions were stopped by adding EDTA and the proteins were digested by adding Proteinase K (New England Biolabs) at 37℃ for 20min. Then RNA duplex and unwound products were separated on native 15% polyacrylamide gel at 90V in 1×TBE buffer at RT. Fluorensent Cy3 signals were detected by BioRad imager.

### CLIP sequencing to identify GLH-1 and GLH-4 targets

CLIP assays were modified from previously described CLIP methods (Chi et al., 2009, Norstrand et al., 2016). Two independent biological replicates were performed for each CLIP assay. Control libraries were also constructed from untagged strains expressing each glh-1 allele and were used to account for unavoidable background RNA binding to the IP matrix.

Independent synchronized populations of young adult animals were exposed to UV light and then subjected to extraction, partial RNase digestion, and FLAG protein immunoprecipitation. Electrophoresis on a denaturing poly-acrylamide gel was then used to separate the proteins in the IP complex, followed by excision of a gel-slice corresponding in size to the GLH protein (and any associated cross-linked RNA fragments). RNA was then eluted from the gel slice and subjected to library construction and deep sequencing.

### Data analysis of CLIP

Raw CLIP sequencing data were first trimmed to remove 3’ adaptors and minimal 15nt reads were kept for later analysis. STAR genome aligner was used for mapping CLIP sequences. Total number of CLIP reads in aggregates were calculated using Htseq and normalized by total read counts. GLH targets were scored by implementing filters of at-least two-fold enrichment over untagged inputs and minimal 5 RPM. Metagene analysis of CLIP data from 5’ to 3’ transcripts was performed by Deeptools using WS262 genome as reference.

## Supporting information

Strains

Oligos

## Acknowledgement

We thank R. Davis, W. Theurkauf and S. Ryder for discussion and suggestions;. D. Conte for reviewing and editing the manuscript; K. Bennent for providing GLH antibodies and K. Ghanta for providing CRISPR RNP method support and strain generation. This work was supported by NIH funding (GM058800 and HD078253) to C.C.M and National Natural Science Foundation of China (NSFC32070628) to E.Z.S. C.C.M is a C.C.M. is a Howard Hughes Medical Institute Investigator.

## Contributions

Conception: S.D, E.S and C.C.M.; Supervision: E.S and C.C.M.; Investigation: S.D, E.S.; Genetics and deep sequencing: S.D, E.S.; Biochemistry: S.D, X.T, T.I.; Bioinformatics: S.D, L.L, A.O.; Writing: S.D, C.C.M.; Review and editing: C.C.M, E.S, D.C.

