## Supplemental Figures for "A family of *C. elegans* VASA homologs control Argonaute pathway specificity and promote transgenerational silencing"

**Figure S1: Purification of PRG-1 and WAGO-1 associated proteins. Fertility of glh-1 mutant strains. *glh* mutants exhibit cold-sensitive RNAi phenotypes.**

(A): Identification of PRG-1 and WAGO-1 associated proteins: 3XFLAG-tagged PRG-1 and WAGO-1 was immuno-precipitated and resolved with SDS-PAGE. Proteins were visualized with silver staining. Red asterisks indicate the position of PRG-1 and WAGO-1.

(B, C): Brood sizes (B) and precent of viable embryos (C) for each indicated strain.

(D) *pos-1* RNAi assay for wild-type, *glh-1(ok439), glh-1(Null) and glh-1(K391A)* animals. Worms are fed with *pos-1* RNAi from L1 stage. Over 300 progenies of each strain were scored for embryonic viability.

(E, F) *Pos-1* RNAi assay for auxin-depleted GLH-1 or *glh-4* null animals double with *prg-1* mutant at different temperature. Worms were fed with pos-1 RNAi food on plates containing both auxin and IPTG.

**Figure S2: GLH-1 and GLH-4 exhibit distinct expression patterns.**

(A, B) Fluorescence microscopy showing GLH-1 and GLH-4 localization within whole gonad. Magnification: 400X. Scale bar represents 20um.

**Figure S3: Abundance of various classes of small RNAs in *glh-1* mutant animals.**

(A) Percent of reads mapped to annotated miRNAs out of total small-RNA sequencing reads.

(B) Scatterplot comparing the numbers of small-RNAs for WAGO targets (1118 genes), CSR-1 targets (3206 genes) and abundant piRNAs (621 genes) between two biological replicates cloned from N2 worms.

(C) Scatterplot comparing the numbers of abundant piRNAs between two biological replicates and cloned from wild-type, *glh-1(ne4712K391A)*, *glh-1(ne4715DQAD)* and *glh-1(Null))* animals.

(D) Scatterplot comparing the numbers of small-RNAs for other categories between two biological replicates and cloned from wild-type, *glh-1(ne4712K391A)*, *glh-1(ne4715DQAD)* and *glh-1(Null))* animals.

(E) Abundance of different classes of small-RNAs (22G-RNAs on WAGO targets, 22G-RNAs on CSR-1 targets and piRNAs) over the time with *de novo* expression of GLH-1(DQAD).

**Figure S4: Ectopic WAGO 22G-RNAs in *glh-1* mutant are suppressed in other *glh* mutants and *prg-1* mutant.**

(A): Box plots showing fold changes on each gene with elevated levels of 22-RNAs when comparing *glh-1(Null)* with WT and *glh-1(Null) prg-1* with WT.

(B): Genome browser view of 22G-RNAs on mog-4 genes cloned from WT, *glh-1, prg-1* and *glh-1;degron::prg-1* double mutant animals.

(C): 171 CSR-1 targets with enhanced WAGO 22G-RNAs. Scatterplot depicting 22G-RNAs cloned via WAGO-1, WAGO-9 and CSR-1 IP from WT and *glh-1 null* animals on 171 targets.

(D): Scatterplots showing 22G-RNAs cloned from WT and *glh-2, glh-3, glh-4* and *glh-1(K391A)* animals. Dotted lines indicate two-fold threshold.

(E): Venn diagram depicting the numbers of CSR-1 targets with marked reduction of ectopic 22G-RNAs in *glh-1* double mutants. *degron::glh-2;glh-1* (orange), *degron::glh-3;glh-1* (green), *degron::glh-4;glh-1* (blue).

(G): Venn diagram depicting the numbers of genes (all) with more than 2-fold reduced 22G-RNAs in *glh-1* K391A (red) and pairwise glh-1 paralog double mutants: glh-1;glh-2 (red), glh-1;glh-3 (orange) and glh-1;glh-4 (blue).

**Figure S5: K391A enhances binding of GLH-1 to PRG-1 and WAGO-1.**

(A): Analysis of IP-MS results by immuno-precipitating GLH-1 WT and GLH-1 K391A protein. Peptide spectra counts for selected co-factors were shown in the table.

(B): A reciprocal Co-IP experiment showing interactions between GLH-1 and PRG-1 or GLH-4 in WT or mutant animals expressing GLH-1(K391A) protein. Presence (+) or absence (-) of RNase I were indicated.

(C): A reciprocal Co-IP experiment showing interactions between FLAG::WAGO-1 and V5::GLH-1 WT or V5::GLH-1 K391A. Presence (+) or absence (-) of RNase I were indicated.

(D, E, F): 280/260 UV absorbance graph was shown to indicate the amounts of proteins of different molecular weights separated by size exclusion chromatography. The collected fractions were resolved by SDS-PAGE and stained with Coomassie Blue.

**Figure S6: GLH-1-associated RNAs are predominantly mRNAs. Enhanced binding of GLH-4 to mRNAs in *glh-1* null mutant correlate with ectoptic WAGO 22G-RNAs. GLH-1 and GLH-1 DQAD enriched on diferent targets regarding mRNA abundance of these targets.**

(A): Table depicting fractions of CLIP reads mapped to each RNA class. (mRNAs, piRNAs, ribosomal RNAs, micro RNAs,  transfer RNAs and other categories.)

(B): metagene analysis of GLH-4 CLIP in WT and *glh-1* mutants over control gene regions and gene regions with ectopic 22G-RNAs.

### Figure S1

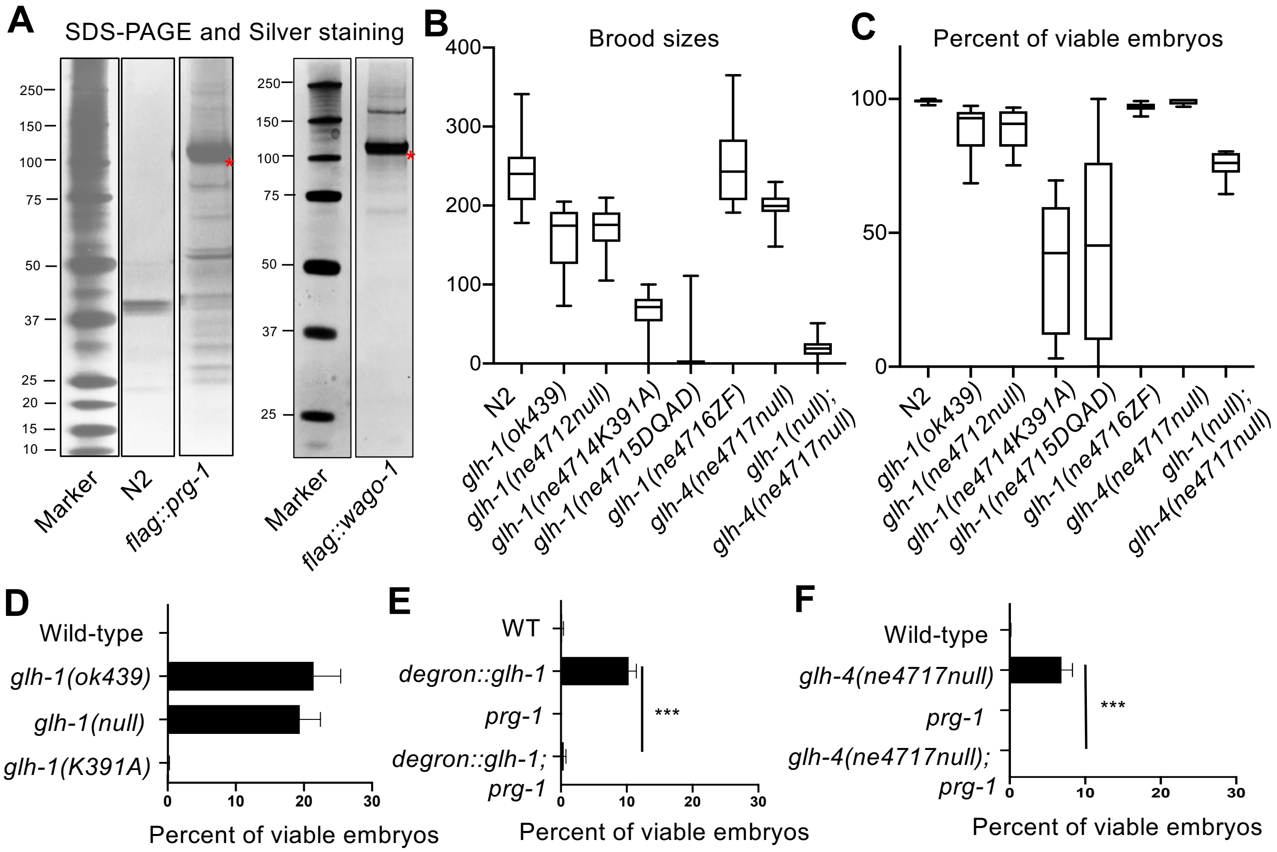
A
SDS-PAGE and Silver staining
250
150
100
75
50
37
25
20
15
10
Marker
N2
flag::prg-1
Marker
flag::wago-1
B
Brood sizes
400
300
200
100
0
N2
glh-1(ok439)
glh-1(ne4712null)
glh-1(ne4714K391A)
glh-1(ne4715DQAD)
glh-1(ne4716ZF)
glh-4(ne4717null)
glh-1(null);
glh-4(ne4717null)
C
Percent of viable embryos
100
50
0
D
Wild-type
glh-1(ok439)
glh-1(null)
glh-1(K391A)
0
10
20
30
Percent of viable embryos
E
WT
degron::glh-1
prg-1
degron::glh-1;
prg-1
***
F
Wild-type
glh-4(ne4717null)
prg-1
glh-4(ne4717null);
prg-1

### Figure S2

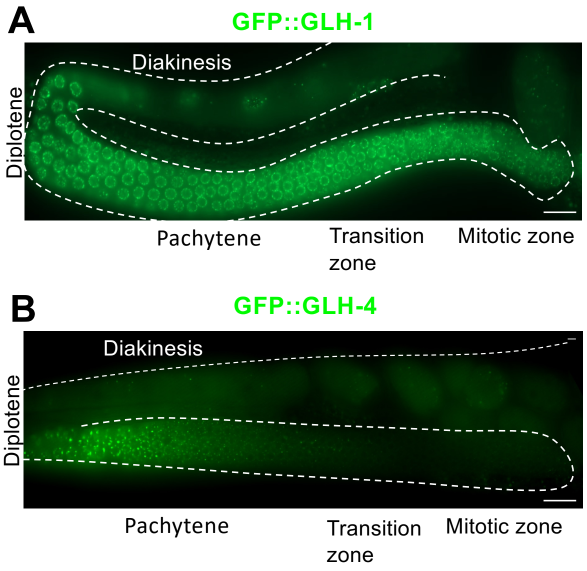
A
GFP::GLH-1
Diakinesis
Diplotene
Pachytene
Transition zone
Mitotic zone
B
GFP::GLH-4
Diakinesis
Diplotene
Pachytene
Transition zone
Mitotic zone

### Figure S3

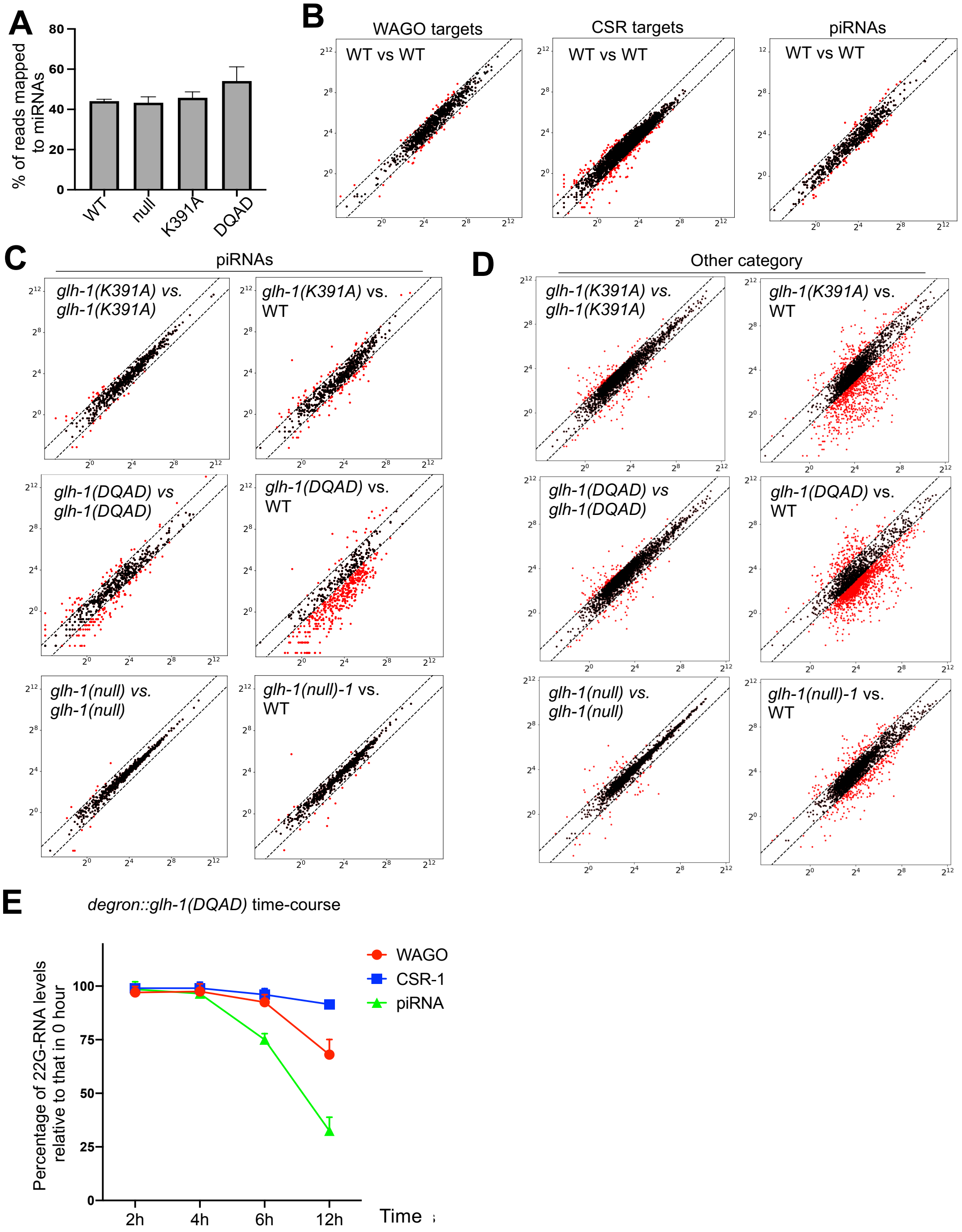

A
% of reads mapped to miRNAs
0
20
40
60
80
WT
null
K391A
DQAD
B
WAGO targets
CSR targets
piRNAs
WT vs WT
C
piRNAs
glh-1(K391A) vs. glh-1(K391A)
glh-1(K391A) vs. WT
glh-1(DQAD) vs glh-1(DQAD)
glh-1(DQAD) vs. WT
glh-1(null) vs. glh-1(null)
glh-1(null)-1 vs. WT
D
Other category
E
degron::glh-1(DQAD) time-course
Percentage of 22G-RNA levels relative to that in 0 hour
0
25
50
75
100
2h
4h
6h
12h
Time
WAGO
CSR-1
piRNA

### Figure S4

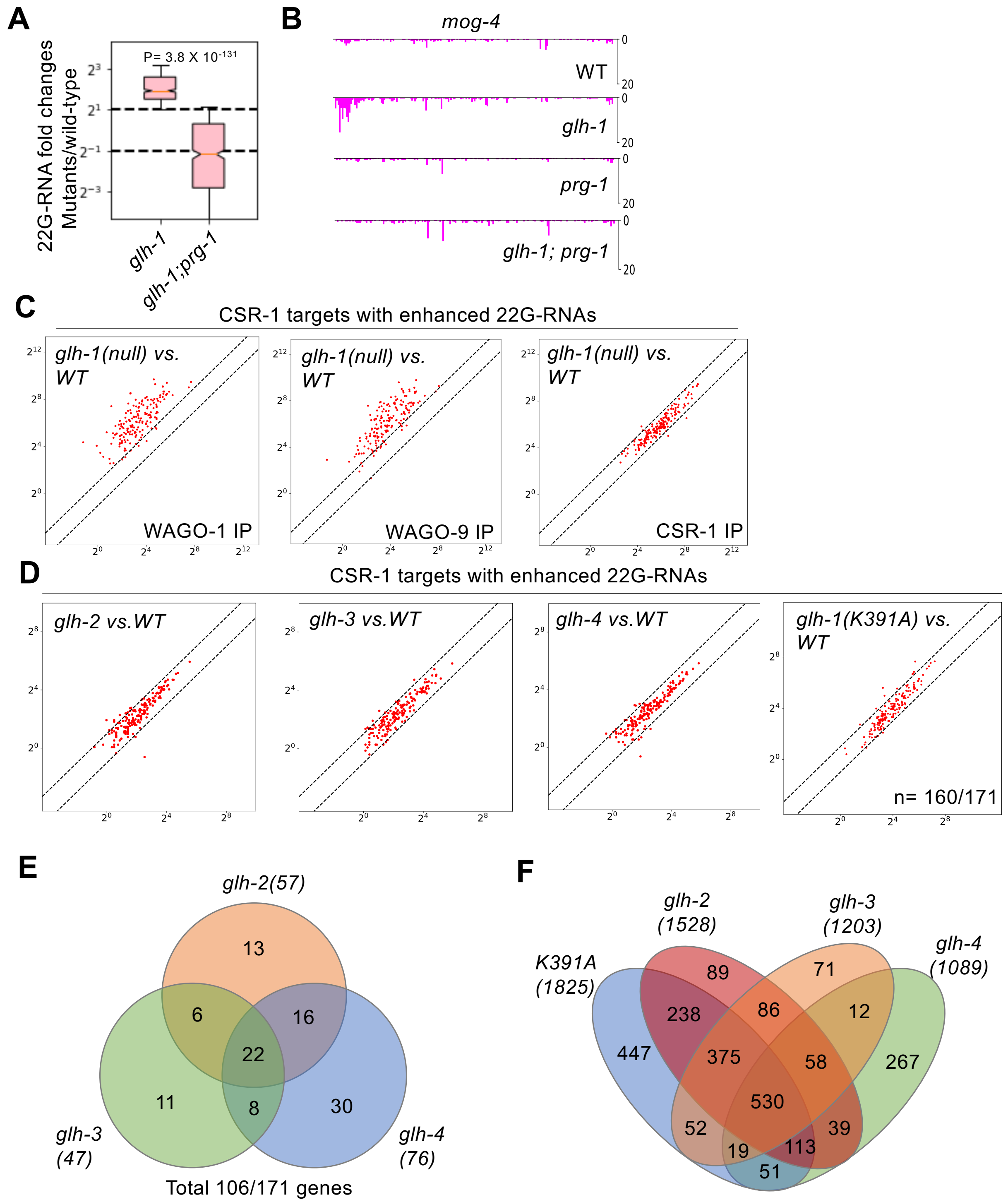
A
P= 3.8 X 10-131
22G-RNA fold changes Mutants/wild-type
glh-1
glh-1;prg-1
B
mog-4
WT
glh-1
prg-1
glh-1; prg-1
C
CSR-1 targets with enhanced 22G-RNAs
glh-1(null) vs. WT
WAGO-1 IP
WAGO-9 IP
CSR-1 IP
D
CSR-1 targets with enhanced 22G-RNAs
glh-2 vs.WT
glh-3 vs.WT
glh-4 vs.WT
glh-1(K391A) vs. WT
n= 160/171
E
glh-2(57)
glh-3 (47)
glh-4 (76)
13
6
16
22
11
8
30
Total 106/171 genes
F
K391A (1825)
glh-2 (1528)
glh-3 (1203)
glh-4 (1089)
89
71
238
86
12
447
375
58
267
530
52
39
19
113
51

### Figure S5

**A**

| Proteins | # peptides by IP-MS | | |
|---|---|---|---|
| | Control | FLAG::GLH-1 IP | FLAG::GLH-1(K391A) |
| GLH-1 | 2 | 2706 | 2346 |
| GLH-2 | 0 | 1116 | 1149 |
| GLH-3 | 0 | 115 | 110 |
| GLH-4 | 0 | 24 | 0 |
| PRG-1 | 0 | 107 | 153 |
| WAGO-1 | 0 | 91 | 164 |
| WAGO-3 | 0 | 42 | 75 |
| WAGO-4 | 0 | 82 | 45 |
| WAGO-5 | 0 | 16 | 15 |
| CSR-1 | 0 | 9 | 10 |
| PAB-1 | 10 | 60 | 128 |
| DEPS-1 | 0 | 45 | 49 |
| ZNFX-1 | 0 | 67 | 83 |
| LAF-1 | 0 | 12 | 0 |
| VBH-1 | 0 | 10 | 0 |
| CGH-1 | 2 | 23 | 0 |

**B**

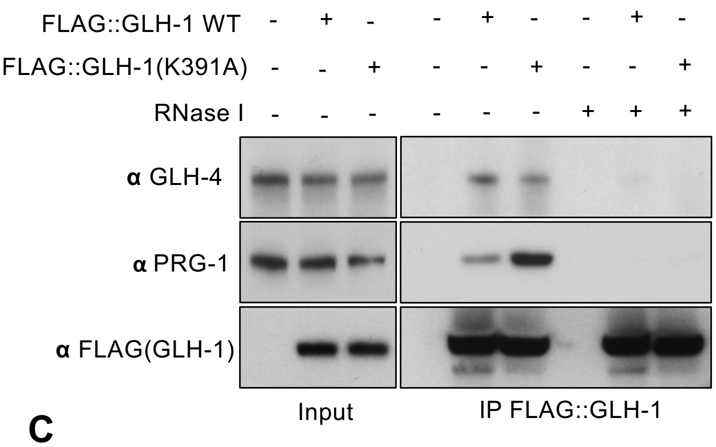


**C**

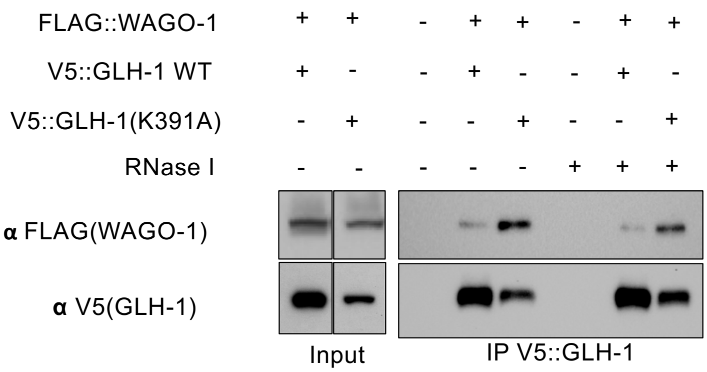


**D**

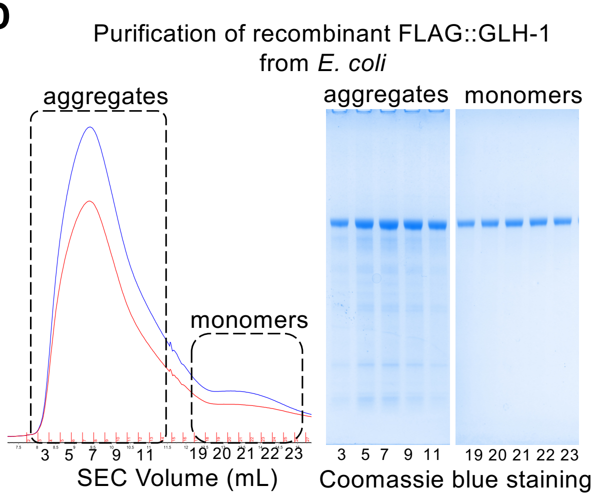


**E**

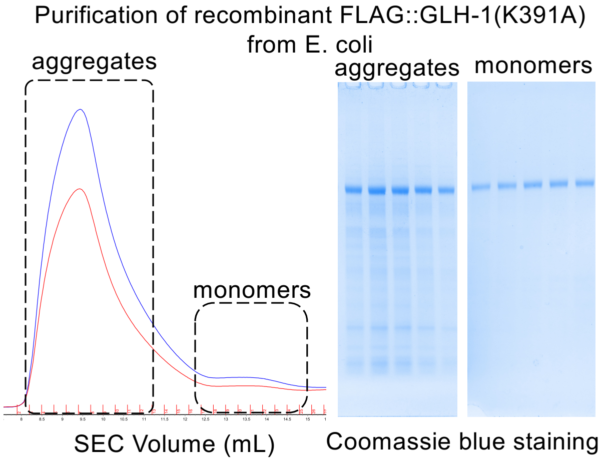


**F**

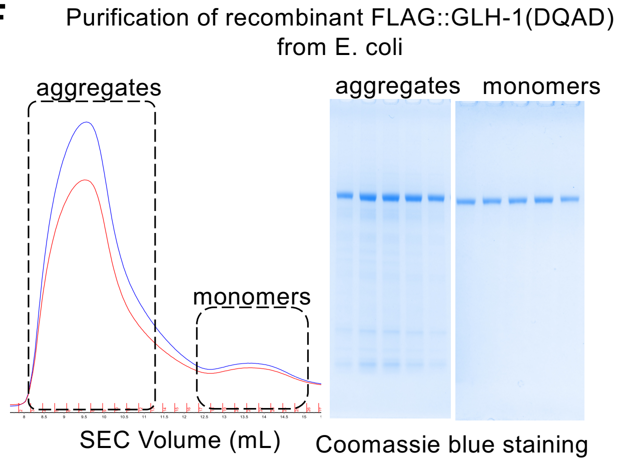

### Figure S6

**A**

Breakdown of mapped CLIP reads

| Samples | mRNA | piRNA | Anti-sense | rRNA | miRNA | tRNA | other |
|---|---|---|---|---|---|---|---|
| GLH-1 | 88% | 6% | <1% | 4% | <1% | <1% | 2% |
| GLH-1;*prg-1* | 89% | <1% | <1% | 5% | <1% | <1% | 4% |
| GLH-1;*rde-3* | 87% | 4% | <1% | 4% | <1% | <1% | 5% |
| GLH-4 | 53% | 25% | <1% | 18% | <1% | <1% | 4% |
| GLH-4 ;*glh-1* | 54% | 21% | <1% | 16% | <1% | <1% | 7% |
| K391A | 88% | 3% | <1% | 2% | <1% | <1% | 6% |
| DQAD 0h | 42% | <1% | <1% | 43% | <1% | <1% | 14% |
| DQAD 2h | 79% | <1% | <1% | 6% | <1% | <1% | 15% |
| DQAD 4h | 49% | <1% | <1% | 35% | <1% | <1% | 15% |

**B**

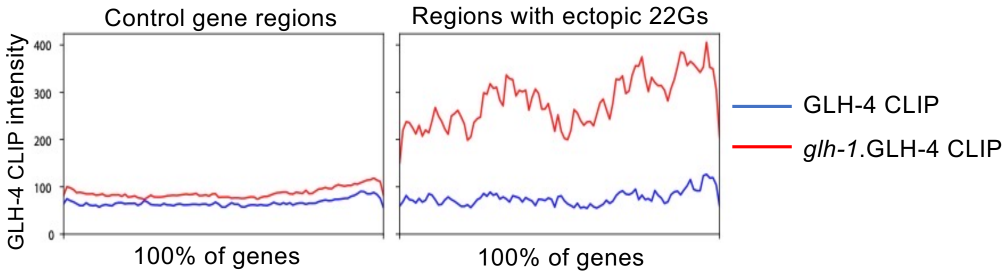
