## Supplementary material for "A family of *C. elegans* VASA homologs control Argonaute pathway specificity and promote transgenerational silencing": Strains

| Strain name: | Genotype | Methods/reference |
| --- | --- | --- |
| N2 | Wild-type | CGC |
| WM604 | *prg-1(ne4484[flag::tev::prg-1]) I* | CRISPR. Ishidate et al., 2018 |
| WM603 | *wago-1(ne4585[3Xflag::TEV::SNAP::WAGO-1])I* | CRISPR Ishidate et al., 2018 |
| WM531 | *neSi12[cdk-1::gfp; Cbr-unc-119(+)] II; unc-119(ed9) III*; *[21ux-1 (anti-gfp) (ne4561)] X* | MosSCI, CRISPR, Shen et al., 2018 |
| WM241 | *neSi11[gfp::cdk-1(RNAe), cb-unc-119(+)] II; unc-119(ed3) III* | MosSCI, Shirayama et al., 2012 |
| WM791 | *glh-1(ok439) I; neSi11 [gfp::cdk-1(RNAe), cb-unc-119(+)] II; unc-119(ed3) III* | Cross |
| WM792 | *glh-1(ne4712[null]) I;neSi11 [gfp::cdk-1(RNAe), cb-unc-119(+)] II; unc-119(ed3) III* | CRISPR |
| WM793 | *glh-1(ne4714[K391A]) I*; *neSi11 [gfp::cdk-1(RNAe), cb-unc-119(+)] II; unc-119(ed3) III* | CRISPR |
| WM794 | *glh-1(ne4715[DQAD]) I*; *neSi11 [gfp::cdk-1(RNAe), cb-unc-119(+)] II; unc-119(ed3) III* | CRISPR |
| WM795 | *glh-1(ne4716[Zinc finger deletion]) I; neSi11 [gfp::cdk-1(RNAe), cb-unc-119(+)] II; unc-119(ed3) III* | CRSIPR |
| WM796 | *glh-4(ne4717[null]) I; neSi11 [gfp::cdk-1(RNAe), cb-unc-119(+)] II; unc-119(ed3) III* | CSRIPR |
| *WM242* | *neSi12[cdk-1::gfp, cb-unc-119(+)] II; unc-119(ed3) III.* | MosSCI |
| *EZS314* | *glh-1(ne4891[null]) I; neSi12 [cdk-1::gfp, cb-unc-119(+)] II; unc-119(ed3) III* | CSRIPR |
| *EZS176* | *glh-1(ne4892[K391A]) I; neSi12 [cdk-1::gfp, cb-unc-119(+)] II; unc-119(ed3) III* | CRSIPR |
| WM825 | *glh-1(ne4919[degron::glh-1]) I; neSi81(Psun-1::tir-1::mRuby::eft 3'utr) IV* | CRSIPR |
| WM826 | *glh-1(ne4919[degron::glh-1]) I; prg-1(ne4920); neSi81(Psun-1::tir-1::mRuby::eft 3'utr) IV* | CRSIPR |
| WM161 | *prg-1(tm872) I;* | ENU |
| WM797 | *glh-4(ne4717[null]) I; prg-1(tm872) I* | Cross |
| WM704 | *glh-1(ne4816[gfp::glh-1]) I* | CRISPR |
| WM798 | *glh-1(ne4895[gfp::glh-1, K391A]) I* | CRISPR |
| WM799 | *glh-1(ne4896[gfp::glh-1, DQAD]) I* | CRISPR |
| WM801 | *glh-1(ne4898[null]) I;; mcherry::pgl-1 I* | CRSIPR |
| WM802 | *glh-1(ne4899[gfp::glh-1, K391A]) I; mcherry::pgl-1 I* | CRSIPR |
| WM803 | *glh-1(ne4900[gfp::glh-1, DQAD]) I; mcherry::pgl-1 I* | CRSIPR |
| WM820 | *prg-1(ne4915[gfp::prg-1]) I; csr-1(ne4515[mcherry::csr-1]) IV;* | CRSIPR |
| WM821 | *glh-1(ne4916[null]) I;*  *prg-1(ne4915[gfp::prg-1]) I; csr-1(ne4515[mcherry::csr-1]) IV;* | CRSIPR |
| WM822 | *glh-1(ne4917[gfp::glh-1, K391A]) I; prg-1(ne4915[gfp::prg-1]) I; csr-1(ne4515[mcherry::csr-1]) IV;* | CRSIPR |
| WM823 | *glh-1(ne4918[gfp::glh-1, DQAD]) I; prg-1(ne4915[gfp::prg-1]) I; csr-1(ne4515[mcherry::csr-1]) IV;* | CRSIPR |
| WM607 | *wago-1(ne[gfp::wago-1]) I* | CRISPR, Ghanta et al., 2019 |
| WM811 | *glh-1(ne4908[null]) I; gfp::wago-1 I* | CRSIPR |
| WM813 | *glh-1(ne4910[gfp::glh-1, K391A]) I; gfp::wago-1 I* | CRSIPR |
| WM815 | *glh-1(ne4912[gfp::glh-1, DQAD]) I; gfp::wago-1 I* | CRSIPR |
| WM800 | *glh-4(ne4897[gfp::glh-4]) I* | CRISPR |
| WM812 | *glh-1(ne4909[null]) I; glh-4(ne4897[gfp::glh-4]) I* | CRSIPR |
| WM814 | *glh-1(ne4911[gfp::glh-1, K391A]) I; glh-4(ne4897[gfp::glh-4]) I* | CRSIPR |
| WM816 | *glh-1(ne4913[gfp::glh-1, DQAD]) I; glh-4(ne4897[gfp::glh-4]) I* | CRSIPR |
| WM824 | *glh-1(ne4894[gfp::degron::glh-1, DQAD]) I; neSi81(Psun-1::tir-1::mRuby::eft 3'utr) IV* | CRSIPR |
| WM521 | *wago-9(ne4336[flag::tev::wago-9]) III;* | CRSIPR |
| WM804 | *glh-1(ne4712[null]) I; wago-9(ne[flag::wago-9]) III;* | Cross |
| WM806 | *glh-1(ne4712[null]) I; csr-1(ne4520[flag::csr-1]) IV* | Cross |
| WM805 | *glh-1(ne4901[null]) I; wago-1(ne4585[flag::wago-1]) I;* | CRSIPR |
| EZS292 | *glh-1(ne4712[null]) I; prg-1(ne4921[degron::prg-1]) I; neSi82(Psun-1::tir-1::mRuby::sun-1 3'utr) II* | CRSIPR |
| EZS219 | *glh-2(ne4922[flag::tev::degron::glh-2]) I; neSi82(Psun-1::tir-1::mRuby::sun-1 3'utr) II* | CRSIPR |
| EZS243 | *glh-1(ne4926) I; glh-2(ne4922[flag::tev::degron::glh-2]) I; neSi82(Psun-1::tir-1::mRuby::sun-1 3'utr) II* | CRSIPR |
| EZS244 | *glh-3(ne4923[flag::tev::degron::glh-3]) I; neSi82(Psun-1::tir-1::mRuby::sun-1 3'utr) II* | CRSIPR |
| EZS220 | *glh-1(ne4926) I; glh-3(ne4923[flag::tev::degron::glh-3]) I; neSi82(Psun-1::tir-1::mRuby::sun-1 3'utr) II* | CRSIPR |
| EZS146 | *glh-4(ne4924[flag::tev::degron::glh-4]) I; neSi82(Psun-1::tir-1::mRuby::sun-1 3'utr) II* | CRSIPR |
| EZS149 | *glh-1(ne4926) I; glh-4(ne4924[flag::tev::degron::glh-4]) I; neSi82(Psun-1::tir-1::mRuby::sun-1 3'utr) II* | CRSIPR |
| WM806 | *glh-1(ne4902[K391A]) I; wago-1(ne4585[flag::wago-1]) I;* | CRSIPR |
| WM827 | *wago-1(ne4585[3Xflag::TEV::SNAP::WAGO-1])I ; glh-1(ne4809[V5::aid::glh-1]) I* | CRSIPR, |
| WM828 | *wago-1(ne4585[3Xflag::TEV::SNAP::WAGO-1])I ; glh-1(ne4925[V5::aid::glh-1, K391A]) I* | CRSIPR, |
| WM808 | *glh-1(ne4903[3Xflag::tev::glh-1]) I;* | CRSIPR |
| WM810 | *glh-1(ne4904[3Xflag::tev::glh-1, K391A]) I;* | CRSIPR |
| WM817 | *glh-1(ne4903[3Xflag::tev::glh-1]) I; prg-1(ne4906) I* | CRSIPR |
| WM818 | *glh-1(ne4903[3Xflag::tev::glh-1]) I; rde-3(ne4907) III* | CRSIPR |
| WM809 | *glh-4(ne4905[3Xflag::tev::glh-4]) I* | CRSIPR |
| WM819 | *glh-1(ne4914[null]) I; glh-4(ne4905[3Xflag::tev::glh-4]) I;* | CRSIPR |
